# Development of genetically-encoded fluorescent KSR1-based probes to track ceramides during phagocytosis

**DOI:** 10.1101/2023.09.27.559623

**Authors:** Vladimir Girik, Larissa van Ek, Isabelle Dentand Quadri, Maral Azam, Maria Cruz Cobo, Marion Mandavit, Isabelle Riezman, Howard Riezman, Anne-Claude Gavin, Paula Nunes-Hasler

## Abstract

Ceramides regulate phagocytosis, however their exact function remains poorly understood. Here we sought 1) to develop genetically encoded fluorescent tools for imaging ceramide, and 2) to use them to examine ceramide dynamics during phagocytosis. Fourteen EGFP fusion constructs based on four known ceramide-binding domains were generated and screened. While most constructs localized to the nucleus or cytosol, three based on the CA3 ceramide-binding domain of KSR1 localized to plasma membrane or endolysosomes. C-terminally-tagged CA3 with a vector-based (C-KSR) or glycine-serine linker (C-KSR-GS) responded sensitively and similarly to ceramide depletion and accumulation using a panel of ceramide modifying drugs, whereas N-terminally tagged CA3 (N-KSR) responded differently to a subset of treatments. Lipidomic and liposome microarray analysis suggested that, instead, N-KSR preferentially binds to glucosyl-ceramide. Additionally, the three probes showed distinct dynamics during phagocytosis. Despite partial lysosomal degradation, C-KSR robustly accumulated at the plasma membrane during phagocytosis, whereas N-KSR becomes cytoplasmic at later timepoints. Moreover, weak recruitment of C-KSR-GS to endoplasmic reticulum and phagosomes was enhanced by overexpression of the endoplasmic reticulum proteins STIM1 and Sec22b, and was more salient in dendritic cells. The data suggest these novel probes can be used to analyze sphingolipid dynamics and function in living cells.

## Introduction

Phagocytosis is a fundamental process whereby external foreign particles are internalized by cells, and the resulting membrane-engulfed compartment, called the phagosome, undergoes a series of maturation steps that lead to the formation of the digestive milieu and degradation of the foreign material (Levin et al.). A large body of literature has highlighted the central role of dynamic phosphoinositide lipid remodeling in controlling this process through the recruitment of a variety of effectors including cytoskeletal, motor or vesicular docking proteins that help guide phagosomes to their fate (Freeman and Grinstein; Levin et al.; Yeung and Grinstein). Sphingolipids are a diverse group of lipids crucial for maintaining cellular membrane structure, fluidity and curvature, that have long been recognized to additionally play bioactive signaling roles (Quinville et al.). Ceramides, a central hub of the sphingolipid network (Hannun and Obeid; Hannun and Obeid), have also been implicated in various steps of the phagocytic process. These span from facilitation of receptor clustering and activation (Avota et al.; Bakr et al.), to initial membrane deformation of phagocytic cups (Magenau et al.; Niekamp et al.), to promotion of phago-lysosome fusion (Hinkovska-Galcheva et al.; Schramm et al.; Wu et al.), and inhibition of antigen presentation in dendritic cells (Fuglsang et al.; Sallusto and Nicol). However, compared to phosphoinositides, less is known about the dynamics, regulation and roles of ceramide and its related lipids during this critical immune process (Saharan et al.).

Arguably, part of the challenge may lie in the difficulty tracking these lipids in living cells (Canals et al.). The most reliable methods such as mass spectrometry do not have the spatial resolution required for subcellular tracking (Buchberger et al.), or rely on labor, cost and material intensive organelle isolation that still suffer from contamination of endoplasmic reticulum (ER) membranes due to tightly tethered membrane contact sites (MCS) that may skew lipid content (Scorrano et al.). In addition, tools such as fluorescent sphingolipid analogs are rapidly metabolized and may partition in membranes in a different manner than their natural counterparts (Canals et al.). Moreover, permeabilization steps necessary for access of anti-ceramide antibodies to intracellular membranes such as phagosomes may disrupt natural lipid localization (Bjørnestad and Lund). Therefore, an alternative approach enabling detection of ceramide in living cells with high spatial resolution is desirable. Genetically encoded lipid probes, typically composed of lipid-binding domains coupled to a fluorescent tag, provide an opportunity to investigate subcellular localization, relative cytosolic accessibility, and dynamics in living cells, and have been extensively employed to dissect phospholipid dynamics during phagocytosis (Hammond et al.). A few described lipid probes specific for sphingolipids have been published (Bakrac et al.; Yamaji et al.). However, no fluorescent ceramide biosensor based on isolated ceramide-binding domains have been described so far, although a range of full-length ceramide-binding proteins have been characterized (Bidlingmaier et al.; Bockelmann et al.; Deng et al.; Zhou et al.). Yet, the localization of these proteins, which are all multi-domain proteins, may be governed by cooperative binding events including ceramide but also involving other factors or domains. An *E. coli* purified glutathione-S-transferase-fusion of the CA3 ceramide-binding domain of the pseudokinase KSR1 was used *in vitro* and demonstrated ceramide-binding specificity (Yin et al.), yet fluorescent versions of this construct for expression in mammalian cells have not been published. In the current report, we document the cloning and screening of 14 putative ceramide-binding genetically encoded fluorescent probes. Out of several potential ceramide-binding biosensors, three constructs all bearing the same CA3 domain of the human pseudokinase KSR1 were characterized further. These were named N-KSR, C-KSR and C-KSR-GS according to which terminal (amino, N-, or carboxy, C-) the fluorescent protein was fused to, and the presence of a glycine-serine (GS) rich linker. The three KSR1-based probes responded to treatments modifying ceramide levels in cells, although changes in both the localization and global fluorescence levels differed between the three constructs depending on the stimulus. Immunostaining, liposome microarray, lipidomic analyses together with further testing of the cellular behavior of the probes indicated that: N-KSR may instead be more sensitive to glycosylated ceramides, C-KSR is less stable than the other two probes, and that both of these factors contributed to the differences in cellular responses observed. Interestingly, differential localization between the three probes, depending on cell type and expression of ER-localized constructs, were documented to occur during phagocytosis. Together our data indicate that these constructs may be useful tools for dissecting subcellular sphingolipid dynamics and warrant further characterization.

## Results

### KSR1-based probes associate to cellular membranes

In previous studies, ceramide has been reported to be present in multiple organelles, including plasma membrane, ER, Golgi, mitochondria, lysosomes and nucleus (Adachi et al.; Götz et al.; Krishnamurthy et al.; Ledeen and Wu; Walter et al.; Wang et al.). In artificial membranes, ceramide, a relatively hydrophobic lipid, readily flip-flops across bilayers (Contreras et al.; Pohl et al.), and although mechanisms that actively maintain bilayer asymmetry of ceramides may exist, to our knowledge they have not been described. Thus, the localization pattern of a ceramide-binding probe could conceivably include binding to several different types of intracellular membranes. To identify ceramide-specific biosensors, four domains that have been previously demonstrated to bind ceramides *in vitro* and/or *in vivo* were selected: 1) the earmuff domain (EMD) of the human SET protein [amino acids (aa) 70- 226] (Mukhopadhyay et al.); 2) the N-terminal C1 domain of atypical human protein kinase C zeta (PRKCZ, also called PKCζ), aa 123-193); 3) the C-terminal C20 domain (aa 405-646) of the same protein PRKCZ (Wang et al.; Wang et al.); and 4) the CA3 domain (also called C1 or cysteine-rich domain, aa 317-400) of the human pseudokinase KSR1 (Yin et al.) (Fig 1A). The selected domains were tagged with EGFP at the amino or carboxy terminus (prefixed with N- and C- respectively) and expressed in mouse embryonic fibroblasts (MEFs). Fusion proteins based on SET (Fig 1Bi-ii) and PRKCZ (Fig 1Ci-iv) were primarily localized to the nucleus. A nuclear export signal (NES) was added to C-SET-EMD (Fig 1Biii) and C-PRKCZ-C20 (Fig 1Cv) to release proteins trapped in the nucleus, a strategy previously used for other lipid biosensors (Goulden et al.). C-NES-PRKCZ-C20 and C-NES-SET-EMD were mainly found in the cytosol, though nuclear localization was still noticeable (Fig 1Biii, 1Cv). In contrast, the expression pattern of KSR1-based constructs depended on the position of the fluorescent tag and linker (Fig 1D). N-terminally-tagged KSR1-CA3 domain (N-KSR) was found primarily at the plasma membrane (Fig Di), while the C-terminally tagged probe (C-KSR) localized to large intracellular vesicles (Fig 1Dii). Examination of the vector-based linker in C-KSR revealed the introduction of a degradation signal RLELKLRILQ (Majeski and Fred Dice), and Western blot as well as immunostainings with organelle markers transferrin, Rab7 and LAMP1 confirmed partial targeting to lysosomes and degradation (Fig S1A-C). Thus, a C-terminally tagged construct with an alternate glycine-serine (GS)-rich linker was also generated and abbreviated C-KSR-GS (Fig 1Diii). This construct displayed a localization similar to N-KSR, although the plasma membrane localization was only salient in a subset of cells, and was more stable than C-KSR (Fig 1Di, 1Diii, Fig S1D). To potentially increase the lipid biosensor affinity (Hammond et al.), a construct expressing two KSR1 CA3 domains in tandem fused at the C-terminus to EGFP was also generated (abbreviated 2X-C-KSR). However, the localization appeared similar to that of C-KSR-GS (Fig 1Div). The localization of both N- and C-KSR constructs depended on two conserved cysteines (C359/C362) whose mutation to serine has previously been shown to decrease ceramide binding *in vitro.* This suggests that their subcellular location might indeed involve ceramide binding (Yin et al.) (Fig 1Dv-vi). When expressed in human Hela cells, probes based on KSR1, SET, and PRKCZ probes show a subcellular localization patterns similar to that seen in MEFs (Fig S2) confirming the idea that they can function similarly in different cellular contexts. Since phagosomal membranes are derived from the plasma membrane and eventually fuse with lysosomes (Levin et al.), we selected three of the KSR1-based probes that showed salient association to such cellular membranes for further characterization: N-KSR, C-KSR and C-KSR-GS. Stable MEF cell lines for N-KSR and C-KSR were successfully generated and employed in subsequent experiments, while C-KSR-GS was characterized by transient transfection.

**Figure 1.**
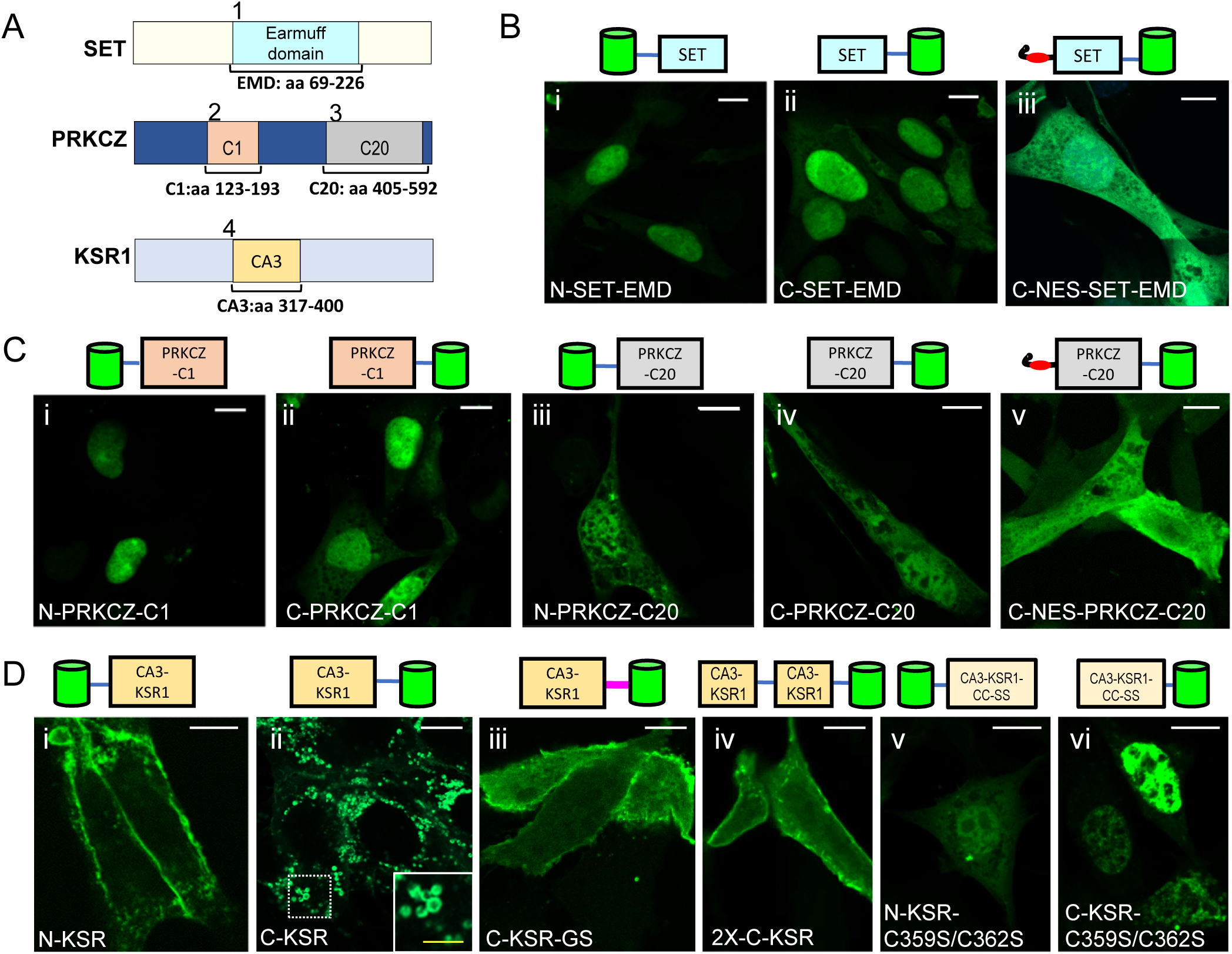
Localization of candidate ceramide probes in mouse embryonic fibroblasts (MEFs). **A)** Schematic representation of ceramide-binding domains used for probe construction from the following source proteins: Human SET earmuff domain (EMD), amino acids (as) 69-226; atypical PKC type zeta C1 domain (PRKCZ-C1, aa 123-193) and C20 domain (PRKCZ-C20, aa 405-592); and KSR1 CA3 domain (KSR, aa 317-400). **B-D)** Confocal images of MEF cells transfected with N-terminally (N-) or C-terminally (C-) EGFP-tagged ceramide-binding domains of SET (B), PRKCZ (C) and KSR1 (D). Constructs preceded by a nuclear export signal are labelled NES, the construct bearing a glycine-serine linker is GS, the tandem construct is 2X, and the cysteine mutants are C359S/C362S. The inset in C-KSR highlights enlarged vesicular structures. White bars = 10 µm, yellow bar = 3 µm.

### KSR-based probes respond to sphingolipid depletion

Next, the behavior of the three KSR probes in response to changes in cellular ceramide levels was investigated. To induce ceramide depletion, cells were exposed to well-known inhibitors of sphingolipid biosynthesis: myriocin, a potent inhibitor of serine-palmitoyltransferase (Miyake et al.), the enzyme which catalyzes the first step in sphingolipid biosynthesis; and fumonisin B1, a selective inhibitor of ceramide synthases (Merrill et al.). Upon ceramide depletion, C-KSR-expressing cells demonstrated a drastic 5-fold decrease in the intracellular fluorescence after a 3-day incubation with myriocin (0.5 µM) or fumonisin B1 (2.5 µM), accompanied by some re-localization to the cytosol in a subset of cells (Fig 2A-B). In comparison, only a 2.5-fold decrease in the probe intensity was observed in N-KSR expressing cells after sphingolipid depletion with either drug (Fig 2C-D). Targeted quantitative tandem mass-spectrometry lipidomic analysis (Guri et al.) confirmed that total levels of all major sphingolipid classes measured including ceramides, glycosylated ceramides (comprising glucosyl and galactosyl ceramides of identical mass, collectively termed hexosyl-ceramides) and sphingomyelins, were markedly depleted by myriocin treatment (Fig 2E, see also Supplementary Data S1). In contrast, total levels of phospholipid species phosphatidylcholine (PC), phosphatidylethanolamine (PE), phosphatidylserine (PS) and phosphoinositide (PI) were unchanged (Fig 2E). In addition, myriocin treatment of cells transiently transfected with C-KSR and cytosolic TagRFP did not produce any noticeable effect on TagRFP expression (Fig S3), confirming a specific effect of myriocin on KSR constructs. In contrast, although no signs of toxicity were noted upon transient transfection of C-KSR-GS alone, 3-day treatment with myriocin or fumonisin B1 of these cells was toxic, precluding analysis. However, overnight treatment with 10 μM fumonisin B1 also resulted in a 2-fold decrease in global cellular expression of C-KSR-GS (Fig 2F-G), indicating that similar to both N-KSR and C-KSR, C-KSR-GS responds to sphingolipid depletion.

**Figure 2.**
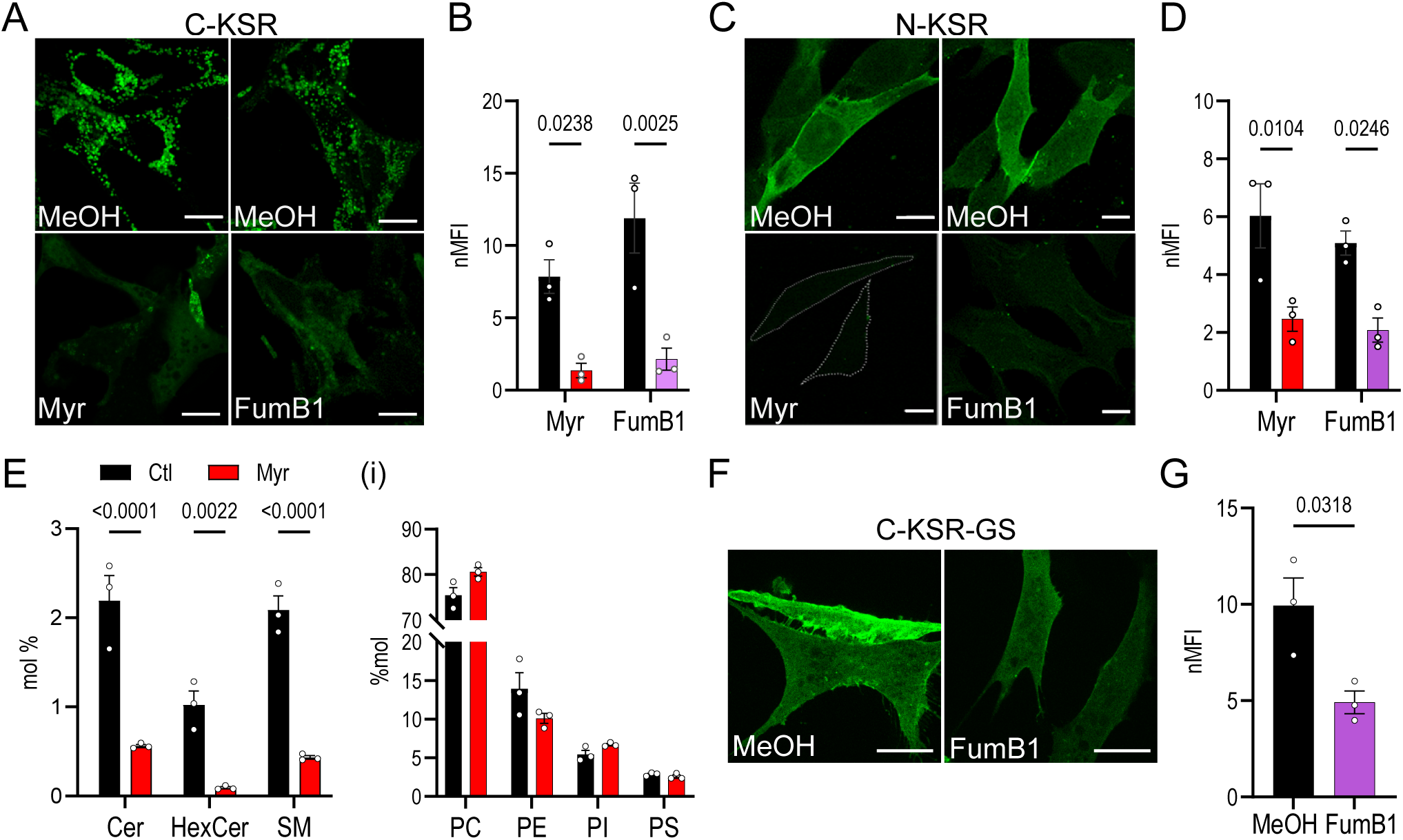
Response of KSR probes to sphingolipid depletion. **A-D)** Confocal images of MEF cells stably expressing C-KSR (A) or N-KSR (B) (in green) treated with; 0.5 µM myriocin (Myr) or 2.5 µM fumonisin B1 (FumB1) FumB1 for 3d; and respective methanol (MeOH) vehicle controls. The quantification of the normalized total cellular mean fluorescence intensity (nMFI) of cells in A,C are shown in B, D respectively. (n= 3 coverslips for each condition). White bars = 10 µm. **E)** Targeted lipidomic analysis of MEF cells stably expressing C-KSR, treated with Myr as above. Left panel shows the quantification of the sum total mass of lipids classified as ceramides (Cer), hexosyl-ceramides (HexCer), and sphingomyelins (SM), normalized to the total mass glycerophospholipids in the same sample (expressed as percent mole, %mol). Inset (i) shows the quantification of total major phospholipid species phosphatidylcholine (PC), phosphatidylethanolamine (PE), phosphoinositide (PI), phosphatidylserine (PS). (n=3 independent biological samples, see also Supplementary Data S1). **F-G)** Similar analysis as in (A-D) of cells transiently transfected with C-KSR-GS (green) and treated overnight with 10 µM FumB1. Bars are means +/− SEM. (n=3 coverslips).

### KSR-based probes react differently to ceramide accumulation

To induce ceramide accumulation at different subcellular locations, two methods were employed. In the first, treatment with the ceramide precursor 16 carbon acyl-chain (C16) fatty acid palmitate for 4h was used to increase ceramide via *de novo* biosynthesis (Kiefer et al.; Tran et al.), which would be expected to increase ceramide in the ER where biosynthesis takes place as well as potentially in the secretory pathway (Hannun and Obeid). In the second, cells were incubated for 30 min with purified bacterial sphingomyelinase from *Staphylococcus aureus* (SMase), which increases plasma membrane ceramides directly by hydrolyzing the choline-ceramide derivative sphingomyelin on the outer leaflet of the plasma membrane, and could potentially increase ceramides additionally within the endocytic system (Abe and Kobayashi; Devaux). Palmitate treatment led to a 3-fold and 1.5-fold increase in global C-KSR and C-KSR-GS signal intensity, respectively, without an obvious shift in subcellular localization (Fig 3A, 3B). This was in contrast to the N-KSR probe which did not show any changes in fluorescence intensity or localization (Fig 3C). Lipidomic analysis confirmed that palmitate treatment induced a 2-fold accumulation of total ceramides, driven by the 4-fold increase in C16-ceramide (Fig 3D, see also Supplementary Data S1). On the other hand, palmitate did not induce changes in total hexosyl-ceramides, nor in total PC, PE and PI (Fig 3D). However, unexpectedly, depletion of the following were noted: total sphingomyelin, driven by decreased C18- and C24-sphingomyelin; C24-hexosyl-ceramide, and total PS (Fig 3D, Di- iii).

**Figure 3.**
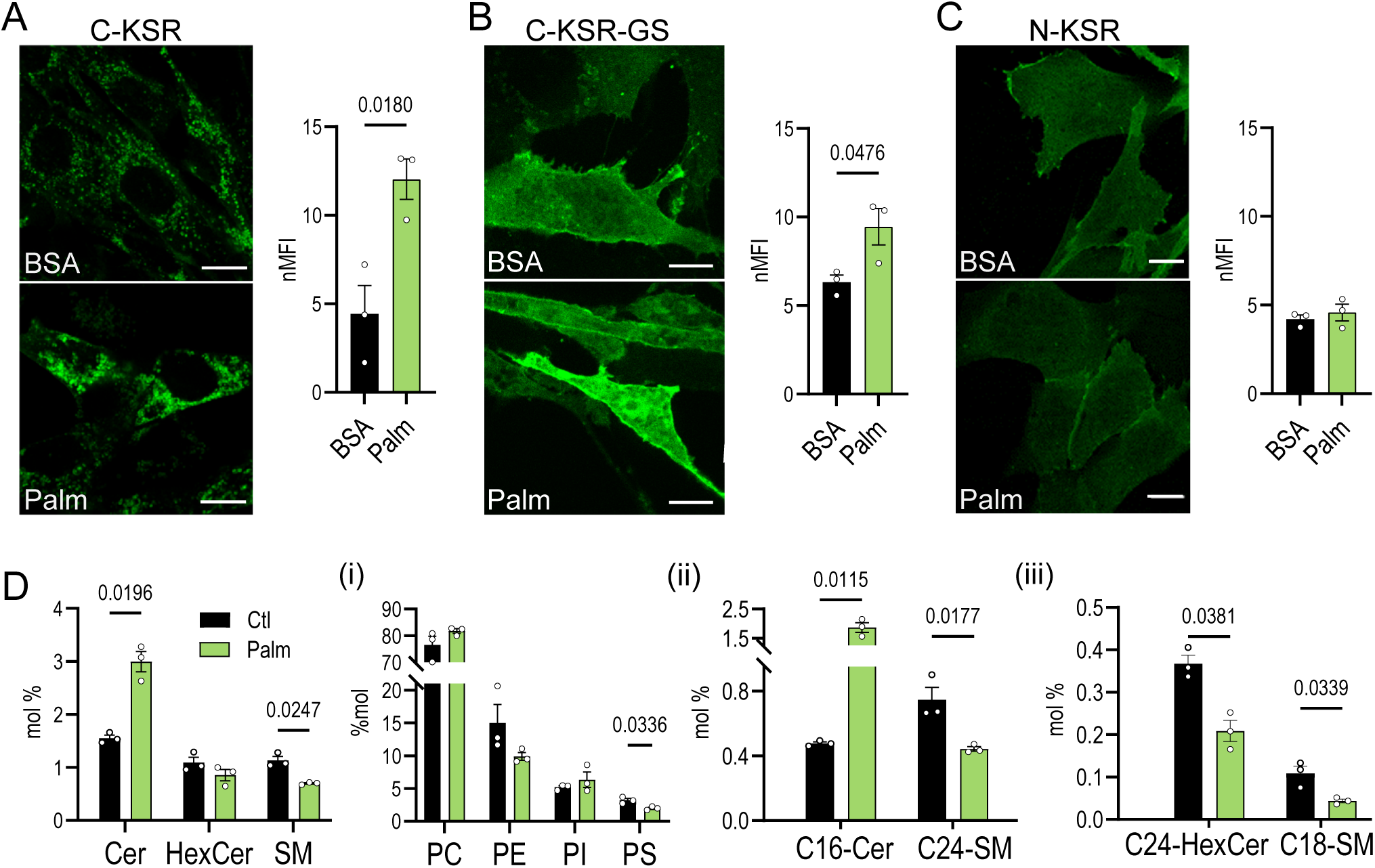
Response of KSR probes to palmitate-induced ceramide accumulation. **A-C)** Confocal images of MEF cells expressing C-KSR (A), C-KSR-GS (B) and N-KSR (C) (in green) treated with; 0.5 mM BSA-palmitate (Palm) or BSA control for 4h. The quantification of the total cellular fluorescence (nMFI) is shown to the right of the respective images (n= 3 coverslips for each condition). **D)** Targeted lipidomic quantification of total Cer, HexCer, and SM in MEF cells expressing C-KSR, treated with Palm as above. Inset (i) shows the quantification of PC, PE, PI and PS. Inset (ii) shows quantification of C16-Cer and C24-SM lipid subspecies. Inset (iii) shows quantification of C24-HexCer and C18-SM lipid subspecies (n=3 independent biological samples, see also Supplementary Data S1). Bars are means +/− SEM. (n=3 coverslips). White bars = 10 µm.

Next, the two C-terminally tagged constructs C-KSR and C-KSR-GS which showed only partial plasma membrane localization under resting conditions were examined for their response to treatment with SMase (0.5 U/ml) for 30 min (Fig 4A). This led both to a small but significant increase in global C-KSR fluorescence intensity (Fig 4B) as well as to a robust 2-fold increase in the recruitment of C-KSR to the plasma membrane (Fig 4A inset, 4C), measured as the fraction of cells displaying plasma membrane localization in SMase treated cells compared to PBS-treated controls (Fig 4C). A significant 1.5-fold increase in the fraction of cells displaying plasma membrane localization of C-KSR-GS was also observed (Fig 4D-E), whereas analysis of N-KSR was not conducted since nearly all cells displayed plasma membrane localization at resting conditions. Lipidomic analysis revealed that SMase induced a dramatic 20-fold accumulation of total ceramide species (Fig 4F), driven by increased C16-ceramide (Fig 4Fii, see also Supplementary Data S1). Total hexosyl-ceramides were not different but an increase in very short-chain C10-hexosyl-ceramides, a lipid species of unknown biological significance, was observed (Fig 4F, Fii). Although total sphingomyelin was not significantly different (Fig 4F), reductions in long-chain C22- and C24-sphingomyelin were observed confirming efficient cleavage of longer-chain species (Fig 4Fii). That cleavage of sphingomyelin was efficient was additionally assessed by staining with a purified EGFP-tagged sphingomyelin-specific probe lysenin (20 µg/ml) (Kwiatkowska et al.), which was lost upon SMase treatment (Fig S4A). Diacylglycerol (DAG) is a molecule with close structural similarity to ceramide that could potentially be indirectly generated by activation of SM synthases upon large increases of ceramide such as those observed under SMase treatment here (Ekman et al.; Goñi and Alonso). Thus, the specificity of C-KSR for ceramide was further investigated by generation of C-KSR-mRFP and co-expression with the DAG probe GFP-PKC-C1(2) (Codazzi et al.). Cells were then treated with either 10 µM ionomycin (Iono) for 10 min, in order to induce plasma membrane DAG accumulation (Botelho et al.) or with SMase as above. Whereas ionomycin treatment but not SMase induced PKC-C1(2) plasma membrane recruitment, the opposite pattern was observed for C-KSR (Fig 4G-H). Treatment with SMase did not affect the localization of PKC-C1(2) when it was expressed alone, nor did it affect the localization of an alternate DAG probe YFP-DBD (Gallegos et al.) (Fig S4C), ruling out potential indirect DAG accumulation after treatment with SMase. Collectively, the data suggest that out of the three probes analyzed, C-KSR is the most sensitive to the changes in cellular levels of ceramides and can also detect locally generated ceramides at the plasma membrane, and that C-KSR-GS responds similarly to C-KSR but with a smaller dynamic range. Most importantly, the lack of response of N-KSR to palmitate suggests a potential difference in lipid binding specificity.

**Figure 4.**
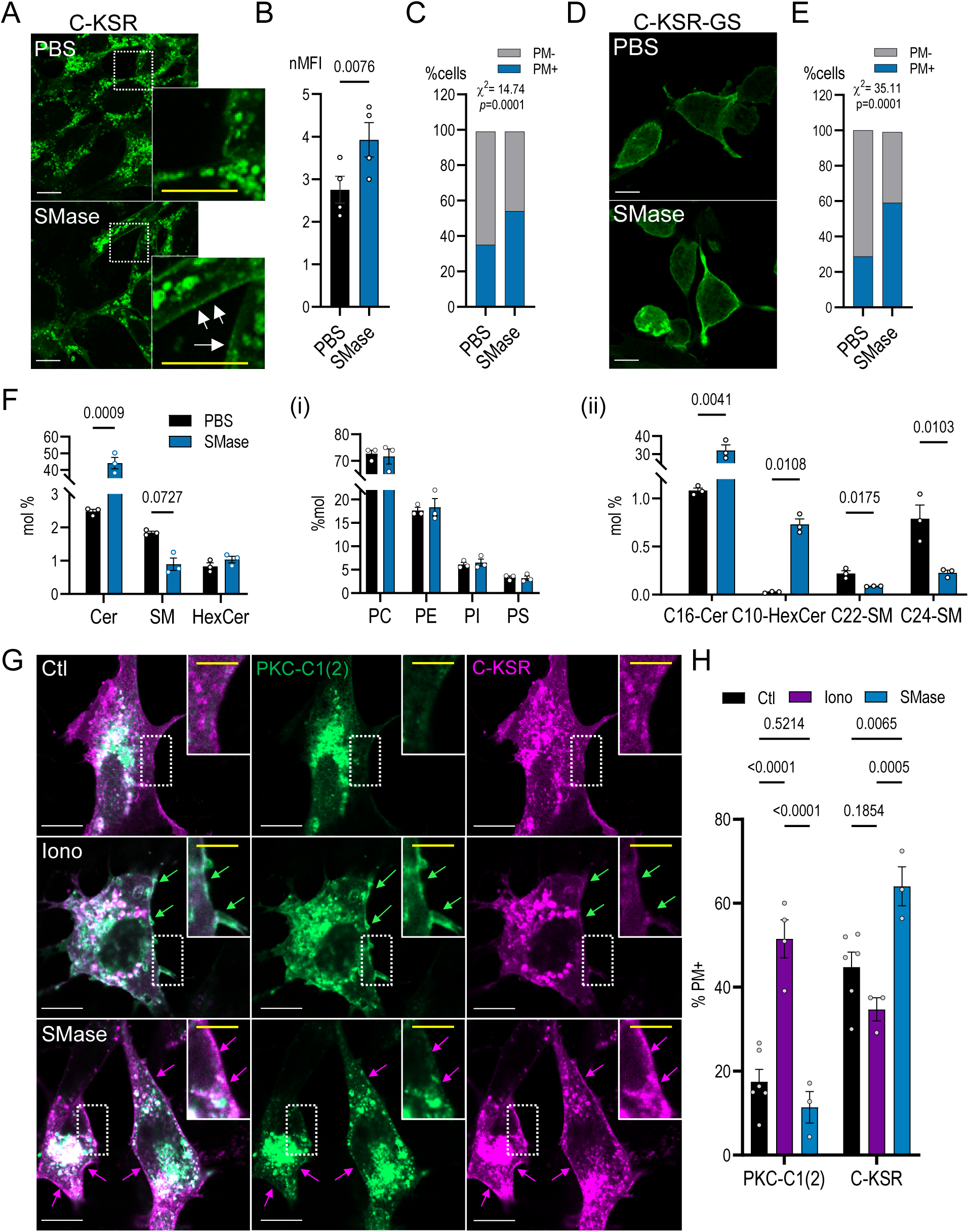
Response of KSR probes to sphingomyelinase-induced ceramide accumulation. **A-E)** Confocal images and analysis of C-KSR (A-C) and C-KSR-GS (D-E) in cells treated for 30 min with 0.5 U/ml sphingomyelinase (SMase). In (B) a quantification of total cellular fluorescence (nMFI) is shown, whereas (C) and (E) show quantification of the fraction of cells displaying plasma membrane localization of C-KSR and C-KSR-GS respectively. (n= 4 coverslips) **F)** Targeted lipidomic quantification of total Cer, HexCer, and SM in MEF cells expressing C-KSR, treated with SMase as above. Inset (i) shows the quantification of PC, PE, PI and PS. Inset (ii) shows quantification of C16-Cer, C10-HexCer, C22-SM and C24-SM lipid subspecies. (n=3 independent biological samples, see also Supplementary Data S1). **G-H)** Confocal images of MEF cells co-transfected with C-KSR-mRFP (magenta) and diacylglycerol (DAG) probe PCK-C1(2)-EGFP (green), control (Ctl, top) or treated with 10 µM ionomycin (Iono) for 10 min (middle), or with SMase as above (bottom). G) Quantification of cells in (G) displaying PM localization of either C-KSR or PKC-C1(2) probe. (n= 6/3/3 coverslips for Ctl/Iono/SMase). Bars are means +/− SEM. White bars = 10 µm, yellow bar = 3 µm.

### N-KSR binds glucosyl-ceramide

To further investigate the mechanism underlying the differences in how the KSR-based probes react to ceramide accumulation, we sought to measure their *in vitro* binding specificity to artificial membranes containing different ceramide species and derivatives. To this end, a liposome microarray-based assay (LiMA) was employed, where arrays of liposomes (small artificial vesicles) of various compositions are deposited inside a microfluidic chip, exposed to purified (or cell lysates containing) fluorescent lipid-binding probes, and binding is assessed by automated fluorescence microscopy (Saliba et al.). To obtain purified probes, the N-KSR and C-KSR probes were re-cloned into super-folding GFP (sfGFP) bacterial expression vectors (Pédelacq et al.) and purified from *E.coli* cultures. Interestingly, significant binding of N-KSR was observed to dioleoyl-phosphatidylcholine (DOPC) liposomes containing 2% glucosyl-ceramide (GlcCer), whereas the trends toward binding to liposomes containing either 10% sphingomyelin or 10% bovine ceramide mixtures were not significant (Fig 5A, Fig S5A, see also Supplementary Data S2). Purified C-KSR tended to aggregate in this assay, and the sample sizes were too small to allow individual statistical analysis of the different membrane types. However, comparison of the binding scores of control liposomes versus those containing 10% ceramide or ceramide derivatives suggests a preference for the latter (Fig 5B). We also studied the binding of C-KSR-GS expressed in lysates from transfected HEK293 cells, to ceramide present in liposomes with either DOPC as the carrier lipid, or in liposomes based on palmitoyl-oleoyl-PC (POPC) that mimic the composition of the inner plasma membrane (IPM liposomes) (Lorent et al.) (Fig 5C, S5C-D). Lysates from HEK cells expressing soluble EGFP and the DAG probe PKC-C1(2) were employed as negative and positive controls, respectively (Fig S5D). Whereas robust and significant binding of PKC-C1(2) to DAG-containing liposomes as compared to IPM liposomes was readily observed (Fig S5D, Supplementary Data S2), the trend for a difference in binding scores of C-KSR-GS to individual ceramide containing liposome compositions as compared to controls (DOPC, POPC or IPM only) alone were not significant (Supplementary Data S2). On the other hand, when pooling liposome data to increase statistical power as above, a comparison of pooled control liposome spots (DOPC, POPC and IPM alone) to DOPC or IPM liposomes containing 5% and 10% ceramides, a small, but significant difference in binding scores was observed in the IPM context (Fig 5C). These data support a preference of C-KSR-GS for ceramide-containing membranes within a more complex lipid context.

**Figure 5.**
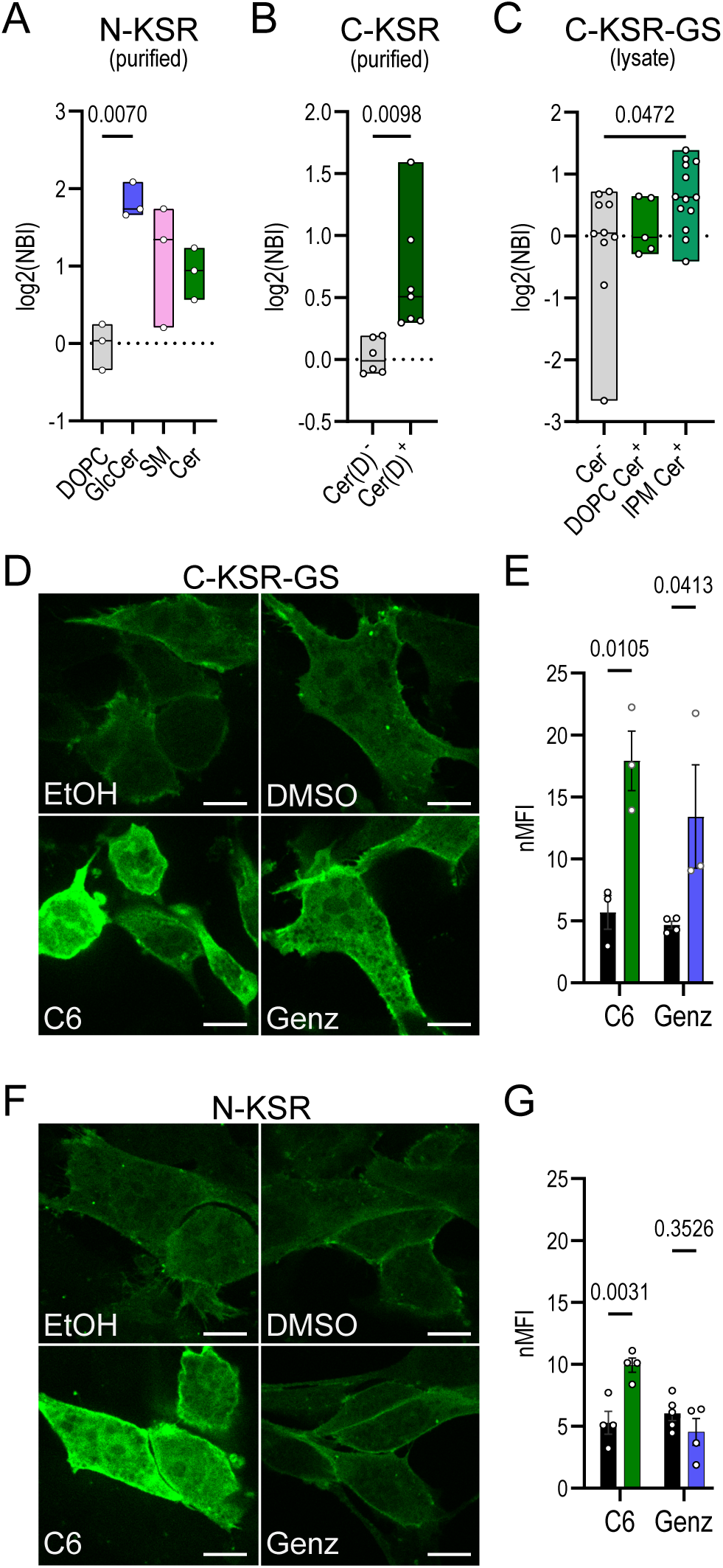
Differential lipid preference of C- and N-tagged KSR probes. **A)** Liposome microarray (LiMA) analysis using purified N-KSR. Binding scores of control liposomes containing dioleoyl- phosphatidylcholine (DOPC), and DOPC liposomes containing 2% glucosyl-ceramide (GlcCer), 10% sphingomyelin (SM) or 10% ceramide mixture (Cer) are shown. Binding scores are expressed as log2 transformed normalized binding intensity (NBI) score ratios to control (n = 3 independent biological samples). **B)** LiMA analysis using purified C-KSR. Pooled binding scores of liposomes either lacking (Cer(D)^-^) or containing ceramide or ceramide derivatives (Cer(D)^+^). Conditions included were: Cer(D)^-^: DOPC alone, and 10%PI(4,5)P2 or 10% sphingosine in DOPC. Cer(D)^+^: 10% ceramide mix (Cer), SM, GlcCer or Cer1P in DOPC. (n=6/7 Cer(D)^-^/Cer(D)^+^ independent biological samples). **C)** LiMA analysis of C-KSR-GS lysates pooled binding scores of control liposomes (Cer^-^) or containing 5% and 10% ceramides in either a DOPC or inner plasma membrane (IPM) mimicking composition (DOPC Cer^+^ and IPM Cer^+^, respectively). Conditions included were: Cer^-^: DOPC, POPC and IPM control liposomes; DOPC Cer^+^: C18-ceramide in DOPC; IPM Cer^+^: ceramide mix and C18-ceramide (n = 10/5/13 Cer^-^/ DOPC Cer^+^/ IPM Cer^+^ independent biological samples). **D-G)** Confocal images of MEFs expressing C-KSR-GS (D) or N-KSR (E) and treated (bottom) with either Cer analog C6-ceramide (200 µM, 2h, left) or 100 µM of the GlcCer synthase inhibitor Genz-123346 (Genz, 1d, right), or their respective vehicle controls ethanol (EtOH) or dimethylsulfoxide (DMSO) (top). (F) and (H) show the quantification of total cellular fluorescence (nMFI) of C-KSR-GS and N-KSR respectively (in F: n=3/3/4/3 coverslips for EtOH/C6/DMSO/Genz; in H: n = 4 coverslips for all conditions). In A-C the box plots show the range (min to max) and the line shows the median value. In E,G, the bars are means +/− SEM. White bars = 10 µm.

Since the *in vitro* conditions tested likely do not fully recapitulate the optimal conditions for lipid binding of our probes, we continued our exploration of the differences between C- and N-terminally tagged probes within the cellular context. Thus, we next examined the response of the two probes displaying the most similar localization pattern (N-KSR and C-KSR-GS) to additional reagents previously shown to increase cellular ceramides. To this end, cells expressing either probe were treated for 2h with 200 μM of the ceramide analogue C6-ceramide (Kjellberg et al.) or overnight with 200 μM of the glucosyl-ceramide synthase inhibitor Genz-123346 (Zhao et al.) or their respective ethanol and methanol vehicle controls. Previously, C6-ceramide has been shown to induce large increases in endogenous ceramides and hexosyl-ceramides, with an only comparatively smaller increase in sphingomyelin (Duran et al.; Kjellberg et al.; Vieu et al.). In contrast, Genz-123346 increased ceramide without affecting sphingomyelin and concomitantly decreasing hexosyl-ceramides (Koike et al.; Zhao et al.). Interestingly, whereas both probes showed a robust increase in cellular fluorescence upon treatment with C6-ceramide (Fig 5E-H), a strikingly different response to Genz-123346 treatment was noted, where an increase in fluorescence was observed with C-KSR-GS but not with N-KSR (Fig 5E-H). Together, these data suggest that a preference of N-KSR for glucosyl-ceramide instead of ceramide could explain the difference in cellular behavior of the N- and C-terminally tagged probes.

### KSR1-based probes behave differently during phagocytosis

In several reports, ceramide generation has been associated with phagocytic ingestion and phagosomal maturation, and shown to be important for pathogen elimination, yet its precise role is still poorly understood (Pathak et al.; Tafesse et al.). Thus, to not only glean some hints at ceramide dynamics (and eventually function) during this important immune process, but also to test our probes under a physiological stimulus, we next investigated the behavior of our three KSR1-based probes during phagocytosis. To this end, we examined MEFs rendered phagocytic by overexpression of the FCGR2A-c-myc. This phagocytic cell model is easier to transfect and manipulate than naturally phagocytic cells, and we and others employing a similar strategy have observed such models recapitulate many aspects of the phagocytic process, including phospholipid dynamics (Bodman-Smith et al.; Criado Santos et al.; Giodini et al.; Hinkovska-Galcheva et al.). Cells expressing each of the three KSR1-based probes (C-, N- and C-KSR-GS) were exposed to 3 µm IgG-opsonized polystyrene beads for 30 min or 135 min, time-points representative of early and late stages of phagosomal maturation (Fig 6A-C). In parallel, sphingolipid levels were measured in C-KSR-expressing cells using targeted lipidomics, where a large accumulation in total cellular ceramides, sphingomyelin and hexosyl-ceramide (4-, 2.6- and 6-fold respectively) was observed at 30 min (Fig 6D, see also Supplementary Data S1). This effect was driven by changes in C16- and C24-ceramide, C24-sphingomyelin, and C16- as well as C24-hexosyl-ceramide species (Fig 6Di-ii). It should be noted that changes in total PC, PI and PS were also observed (Fig 6Diii- iv). Interestingly, 30 min after the incubation with IgG-coupled beads, accumulation of C-KSR at the plasma membrane, similar to what was observed after incubation with bacterial SMase, was noted (Fig 6A, see also Fig 4A). Exploring this plasma membrane localization further, we found that plasma membrane staining of C-KSR could still be detected at 45 min and 60 min after phagocytosis initiation, yet was lost at later stages, after 90 and 135 min (Fig S6A). Similarly, plasma membrane localization of N-KSR and C-KSR-GS was salient at 30 min (Fig 6B-C), even though both constructs were already found at the plasma membrane under basal conditions, whereas at 135 min the global N-KSR signal was not as clearly discerned (Fig 6B). Moreover, whereas N-KSR was completely absent from phagosomes (Fig 6B, insets), in C-KSR-GS-expressing cells, accumulation around phagosomes was detected in a subset of phagosomes (Fig 6C, arrows). In some instances, a pattern of small flattened cisternae or protruding tubular structures could be seen around phagosomes, which were reminiscent of membrane contact sites (MCS) between the phagosomal membrane and ER, structures that have been studied extensively by our laboratory in the past (Criado Santos et al.; Guido et al.; Nunes-Hasler et al.; Nunes et al.) (Fig 6C, arrows). To confirm whether such structures could indeed represent ER, C-KSR-GS was then co-expressed with several ER-targeted constructs, including: a soluble ER-targeted RFP TagRFP-KDEL, a calcium regulator and MCS marker mCh-STIM1 and mCh-Sec22b, a protein regulating traffic between ER and Golgi that we have recently shown to tether STIM1-independent MCS (Criado Santos et al.; Nunes et al.) (Figs 6E-F, Fig S6B). Interestingly, although overlap between C-KSR-GS and KDEL was not easily discerned, likely due to low overall KDEL expression (Fig S6B), overlap between C-KSR-GS and STIM1 as well as C-KSR-GS and Sec22b was observed in some but not all instances of periphagosomal accumulation of the fluorescent proteins (Fig 6E-F). Finally, we expressed the N-KSR and C-KSR-GS probes in a naturally phagocytic dendritic cell line JAWS II. Similar to MEFs, N-KSR did not localize to the phagosomal or other organellar membranes and appeared to be depleted at late phagocytic time points (Fig 7A). In contrast, robust C-KSR-GS accumulation at the phagosomal membrane could be discerned on a subset of phagosomes, as well as on internal structures reminiscent of the ER (Fig 7B). Together these intriguing observations suggest the following: first, that the large changes in ceramide during the first 30 min of phagocytosis detected in whole-cell MEF extracts are likely occurring at the plasma membrane and are best detected by C-KSR which is the most sensitive of the three probes at detecting changes of plasma membrane ceramides, whereas changes in glucosyl-ceramide may be explored instead using the N-KSR probe. Secondly, C-KSR-GS may be more useful to detect ceramide species on intracellular membranes including phagosomal and ER membranes. Finally, the data also hint at dynamic ceramide fluxes, potentially at contact sites between phagosomal membrane and ER.

**Figure 6.**
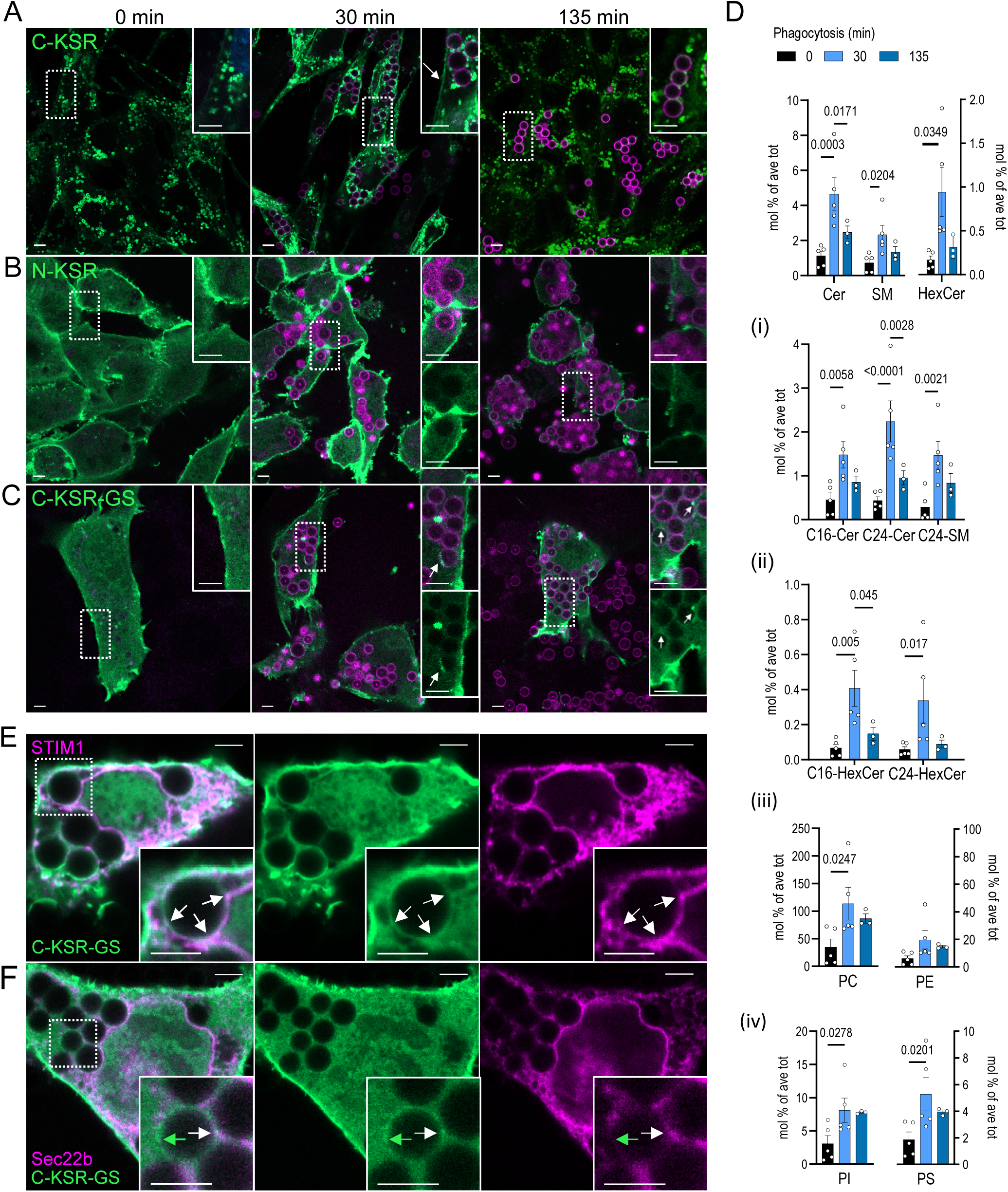
Behavior of KSR probes during phagocytosis in MEFs. **A-C)** Confocal images of phagocytic MEFs expressing (in green) C-KSR (A), N-KSR (B) and C-KSR-GS (C), exposed to IgG-Alexa647-coupled beads at 1:10 cell:bead ratio for 0 (left), 30 (middle) and 135 (right) min. **D)** Targeted lipidomic analysis of MEF cells stably expressing C-KSR, treated exposed to IgG-beads as above. Top panel shows the quantification of the sum total Cer, HexCer, SM. Quantification of the species showing significant changes grouped by fatty acid chain length for Cer and (i) or HexCer (ii). Insets (iii) and (iv) show quantification of total phospholipids for PC and PE in (iii), PI and PS in (iV), (n=3 independent biological samples, see also Supplementary Data S1). **E-F)** Confocal images of phagocytic MEFs co-transfected with C-KSR-GS (green) and membrane contact site proteins mCh-STIM1 (E, magenta) and mCh-Sec22b (F, magenta), and exposed to IgG-beads as above. White arrows show periphagosomal regions where overlap of fluorescent protein signal is discerned, green arrow shows regions with green fluorescence only. Bars are means +/−SEM. White bars = 3 µm.

**Figure 7.**
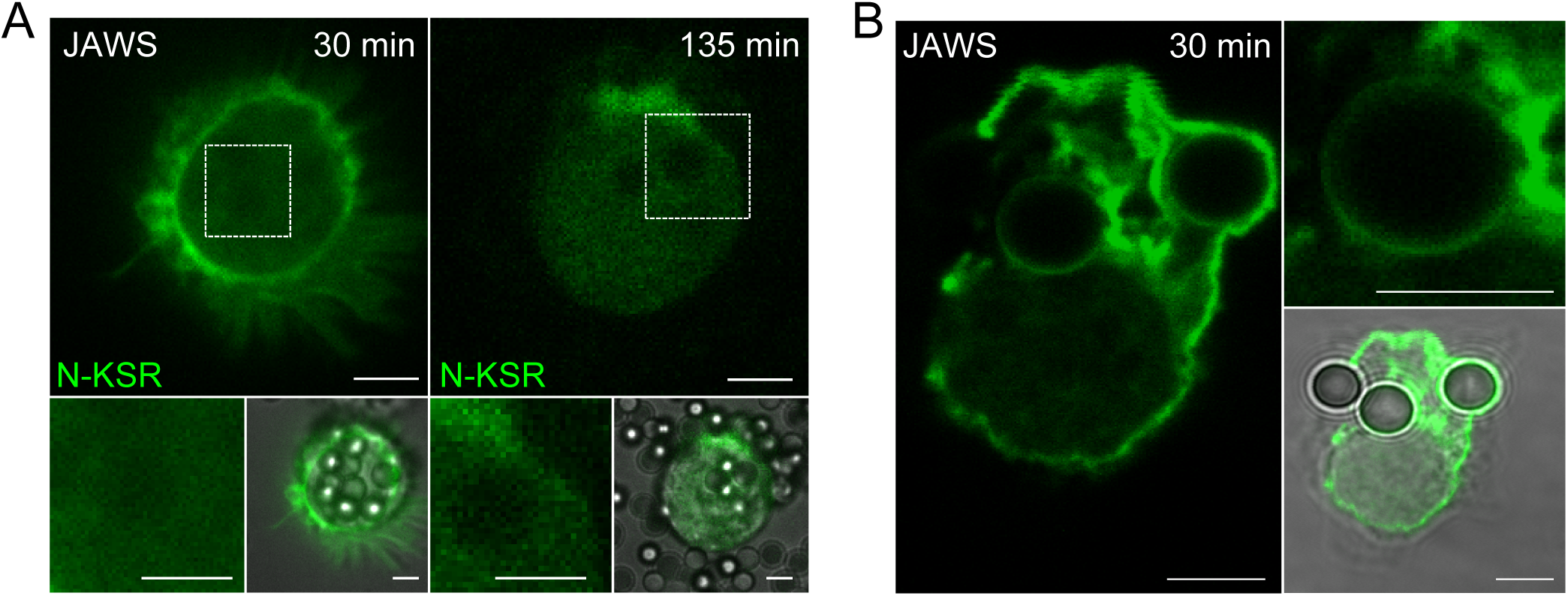
Behavior of KSR probes during phagocytosis in dendritic cells. **A-B)** Confocal images of dendritic cell line JAWS transfected with (in green) N-KSR (A) exposed to IgG-coupled beads at 1:10 cell:bead ratio for 30 (left) and 135 (right) min, and C-KSR-GS (B) exposed to beads for 30 min. Brightfield images are shown to confirm presence of internalized beads. Bars = 3 µm.

## Discussion

Despite several proteins reported as ceramide binders (Bockelmann et al.; Canals et al.; Zhou et al.), to our knowledge no genetically encoded ceramide-specific probes per se have been reported, though whole GFP-tagged ceramide-binding proteins were employed to track subcellular ceramide in a limited number of studies (Canals et al.). Out of 14 candidate biosensor constructs reported here, we focused on three based on the KSR1 CA3 (317-400) ceramide-binding domain which displayed membrane localization when expressed in cells. In contrast, constructs based on PRKCZ and SET were targeted to the nucleus, without obvious association to cellular membranes. Since sphingolipids have been reported to be present in both the nuclear membrane as well as nuclear matrix (Ledeen and Wu), and indeed SET has a defined nuclear activity (Adachi et al.), it is possible that these proteins may detect ceramide in the nuclear matrix rather than simply being misfolded (Albi and Viola Magni). Future studies employing stimuli such as DNA damage, known to change nuclear ceramides may still help to verify the utility of these constructs.

In contrast, though nuclear shuttling has also been reported for KSR1, this scaffolding regulator of MAPK signaling normally resides in the cytosol in resting conditions (Brennan et al.). It is recruited to the plasma membrane upon growth factor stimulation, which induces both changes in phosphorylation of multiple KSR1 residues, as well as the activities of neutral and acidic SMases, generating ceramide (Mathias et al.; Razidlo et al.). Thus KSR1-based probes are particularly poised to detect ceramide in the context of plasma membrane lipids. The baseline localization of N-KSR, 2x-C-KSR and C-KSR-GS, as well as recruitment to the plasma membrane upon SMase treatment of C-KSR and C-KSR-GS are all consistent with this notion. In addition to ceramide binding, mutagenesis studies suggest that phosphorylation of T274 and S392 contribute to KSR1 stability. Notably, dephosphorylation of S392, a residue found within the CA3 domain, participates in membrane re-localization upon growth factor activation (Brennan et al.; Razidlo et al.). Whether S392 in the CA3 of our KSR constructs is constitutively dephosphorylated, or whether separation from other KSR1 domains is sufficient to allow conformational changes favoring ceramide binding remains to be defined. However, full-length KSR1-S392A localizes to the nucleus rather than the plasma membrane (Brennan et al.). Several studies have reported direct ceramide binding of the KSR1 CA3, including *in vitro* reports using an N-terminal GST-fusion with a CA3 of similar composition (aa 320-388) as in our own study (aa 317-400), though others, notably some employing a truncated CA3 (aa 334 to 377), have failed to detect direct binding (Pérez-rivas et al.; Yin et al.; Zhang et al.; Zhou et al.). On the other hand, the structure of this truncated CA3 has been resolved and shows a shallow ceramide binding pocket that was identified based on similarity to other C1 domains, involving residues L342, V345, M353, I354 and F355 (Zhou et al.). Thus, the KSR1 CA3 would not be expected to, on its own, interact with the fatty acid chain or to be able to extract lipids from membranes, but rather to interact with lipid head-groups.

In the characterization of our probes, a surprising observation was that the most salient change in the behavior of the KSR probes in response to ceramide-modifying drugs was a change in total cellular fluorescence. This contrasts with similar phospholipid probes, which usually change their locations rather than their total fluorescence. The effect was consistently observed in both induction and depletion of ceramides and across all drug treatments tested (Figs 2-5). For myriocin treatment, a simple toxicity effect (which often accompanies changing levels of ceramide (Hannun and Obeid) was ruled out since levels of cytosolic RFP were unchanged. In addition, with SMase treatment, neither the two DAG probes reacted in this manner (Fig 4, S4). A potential explanation is that the CA3 domain itself is somewhat unstable and subject to an intrinsic degradation that is mitigated by membrane binding. Indeed, full-length KSR1 binds cytosolic chaperones Hsp70, 90 and 68, (Stewart et al.), and KSR1 is ubiquitinated, though at a site outside the CA3 domain (Matheny et al.; Rinaldi et al.). Moreover, phosphorylation at both T274 and S392 increases the half-life of KSR1 from 1hr to 6hr (Rinaldi et al.). Such a fast turnover time is consistent with our results which were observed even with treatments as short as 30 min. A requirement for turnover time may also explain why a smaller increase in total fluorescence was observed with the shorter SMase treatment, despite the fact that changes in total ceramides were larger. However, whether phosphorylation, ubiquitination, or chaperone binding might also be modified, increasing stability also of the cytosolic fraction of the probe, will be important to define in future studies.

In addition to its potential influence on cellular probe behavior, an intrinsic instability, or requirement for post-translational modification of the CA3 domain to prevent aggregation or precipitation in solution, and increase ceramide binding efficiency, could well explain the weak though detectable *in vitro* binding observed for C-KSR-GS and C-KSR to artificial membranes containing ceramide or ceramide-derivates. Nevertheless, a significant and surprising interaction of the N-KSR probe to glucosyl-ceramide-containing liposomes was observed. Though unexpected, such a modification to the lipid specificity of the CA3 domain is consistent with the difference in behavior between N- and C- constructs to palmitate, Genz- 123346 as well as to phagocytic stimuli. Future studies with drugs or genetic manipulation of enzymes such as glucosyl and galactosyl ceramidases or transporters such as TMEM16F (Suzuki et al.) would be useful to corroborate these finding and determine more precisely the specificity of this probe for glycosylated or potentially phosphorylated ceramides in the cellular context. In addition, it would be worth to explore whether, for example, the creation of S392A mutants of the C-KSR-GS construct, which lacks MEK-interaction domains that leads to nuclear localization (Brennan et al.), may improve probe stability in solution, allowing *in vitro* ceramide-binding to be more robust. Overall, the examination of our three KSR1-based probes under different stimuli (summarized in Fig 8) demonstrated that these probes may each have their own utility as such or serve as a basis for future designs and studies aimed at dissecting ceramide, and potentially glucosyl-ceramide, dynamics in cells. The exacerbated instability of C-KSR, which prevented constitutive plasma membrane localization, proved in fact to be advantageous in the way that it allowed for a greater dynamic range and more salient plasma membrane recruitment with a phagocytic stimulus than C-KSR-GS. Thus, like conditional instability employed in nanobody engineering (Tang et al.), a future design could build upon these observations to include proteasome targeting signals, which may be less disruptive than the current design to membrane homeostasis (Zhao et al.), to obtain a sensitive on-off biosensor for plasma membrane ceramide.

**Figure 8.**
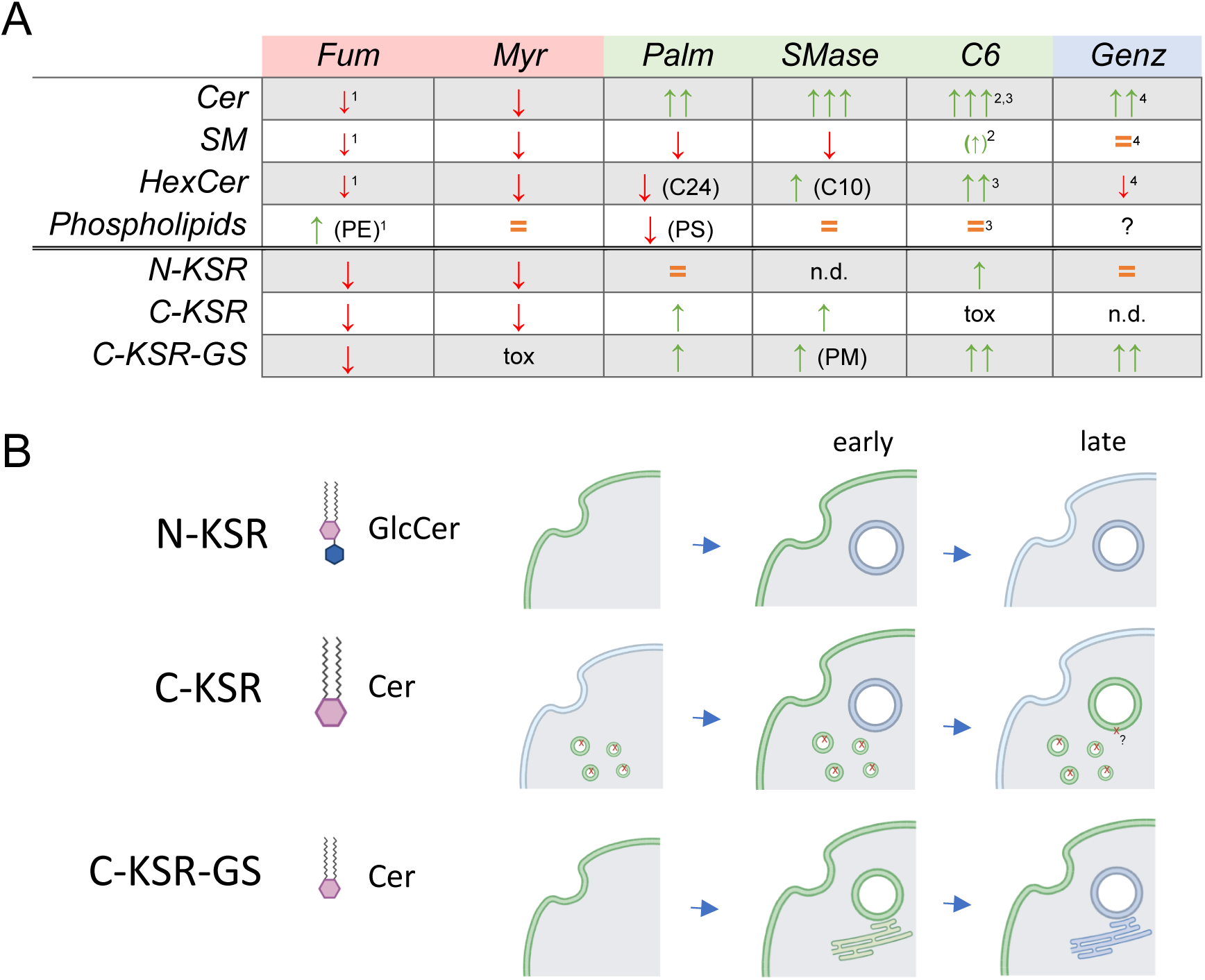
Summary and model. **A)** Table summarizing effect of ceramide-modifying treatments used in this study on major lipid classes (ceramides (Cer), sphingomyelins (SM), hexosyl-ceramides (HexCer) and phospholipids) and on the 3 KSR1-based probes. In cases where only certain sub-species of lipids were affected, the major sub-species affected in listed in brackets. Decreases are shown as a red arrow pointing down, increases as green arrow(s) pointing up, and similar levels as an orange equal sign. Tox = toxicity. n.d. = not determined. PM = plasma membrane. Myr (myriocin), Palm (palmitate) and SMase (sphingomyelinase) effects were measured in this study. The effects of Fum (fumonisin B1), C6 (C6-ceramide) and Genz (Genz-123346) are extrapolated from the literature. (^1^Merrill et al, 2011; ^2^Vieu et al, 2002; ^3^Duran et al. 2012; ^4^Koike et al, 2019). **B)** Hypothetical model of the behavior of N-KSR, C-KSR and C-KSR-GS before and during phagocytosis. N-KSR (top), with a higher sensitivity for glucosyl-ceramides (GlcCer) is lost at later stages of phagocytosis. C-KSR (middle) and C-KSR-GS (bottom) are both more sensitive to ceramides (Cer). Higher dynamic range and turnover of C-KSR may help more sensitive detection of ceramide changes at the plasma membrane during early stages of phagocytosis, but presence in lysosomes and phagosomes may represent a partially degraded probe. The more homogeneous and higher cytosolic expression of C-KSR-GS may make it most suitable for detecting ceramide changes on the cytosolic leaflet of organellar membranes such as the phagosome and endoplasmic reticulum. Created with BioRender.com

Finally, our intriguing observations of the differential behavior of N-KSR and C-KSR-GS during phagocytosis further support the notion that these two very similar probes are responding to different lipid changes in the cell. One important notion to consider is that such cytosolic probes will only detect lipids that are present on the cytosolic leaflet, or when luminal leaflets are exposed to the cytosol for example during plasma membrane or lysosomal disruption. Indeed, the sphingomyelin probe lysenin has recently contributed greatly to our understanding of the role of sphingomyelin (and indirectly ceramide) in membrane repair and antigen cytosol transfer in dendritic cells (Canton et al.; Childs et al.; Niekamp et al.). Though several studies indicate that ceramide readily flip-flops across lipid bilayers (Contreras et al.; Goñi et al.; Pohl et al.), this is not so for complex sphingolipids such as glucosyl-ceramide, where trans-bilayer transporters might maintain or establish gradients in response to cellular signals as occurs for PS (Suzuki et al.; Williamson). Changes such as the loss of the N-KSR signal might equally represent a loss of total glucosyl-ceramide as it might simply reflect its removal from the cytosolic leaflet, which could explain why global changes in hexosyl-ceramides was not detected by lipidomic analysis, though compensating changes in galactosyl-ceramide could also provide an explanation. However, that the exciting ER localization of C-KSR-GS upon certain conditions is simply a result of misfolding is unlikely, and investigation of whether this truly represents ceramide flux is ongoing in our laboratory. Although clearly more work will be required to define the behavior and lipid specificity of these novel biosensors, we believe these constructs represent useful new tools to explore sphingolipid regulation and function in living cells.

## Materials and Methods

### Reagents

The following antibodies (antibody Name/catalog #/dilution) were from: Abcam (USA) – anti-Rab7/ ab137029/1:100; anti-Rab5/ab18211/1:100; Enzo Life Sciences (USA) – anti-LAMP1/ ADI-VAM-EN001/1:50, BD Biosciences (USA) – anti-TGN38/ 610898 /1:200; Thermo - AF555 Goat anti-mouse IgG/ A-21422/1:500, AF555-goat-anti-rabbit IgG/A-21428/1:250; Santa-Cruz Biotechnology - Goat anti-rabbit IgG-HRP/1:5000, BioRad (USA) – Goat anti-mouse IgG-HRP/170-6516/1:3000. All lipid standards for lipidomic analysis were from Avanti (Sigma/Merck) amounts given per 6-cm cell culture dish: DLPC (12:0-12:0 PC, 0.4 nmol/dish, cat. #850335), PE31:1 (17:0-14:1 PE, 1 nmol/dish, cat # LM-1104), PI31:1 (17:0-14:1 PI, 1 nmol/dish, cat # LM-1504), PS31:1 (17:0-14:1 PS, 3.3 nmol/dish, cat# LM-1304), CL56:0 (14:0 Cardiolipin, 0.7 nmol/dish, cat#710332), C12SM (C12 Sphingomyelin d18:1-12:0, 0.1 nmol/dish, cat#860583), C17Cer (C17 Ceramide, d18:1-17:0, 2.5 nmol/dish, cat #860517), C8GC (C8 Glucosyl(B)Ceramide, d18:1-8:0, 0.5 nmol/dish, cat#860540). Lipids employed in the liposome microarray analysis are listed in Supplementary Table S1. Chemicals were purchased from Sigma/Merck unless otherwise stated, (Name/catalog #): HPLC grade Methanol/4860-1L-R, HPLC grade Chloroform/650471, tert-Butyl Methyl Ether (MTBE)/650560, Methylamine/534102; LC-MS grade water/1153331000. Thermo: HPLC grade n-butanol/11378197. Plasmids used and generated in this study are listed in Supplementary Table S2.

### DNA cloning

All newly made constructs were verified by sequencing. Constructs expressing candidate ceramide probes were constructed by subcloning the human KSR1 CA3 domain (aa 317-400, XM_047436985), N-terminal C1 domain of human PRKCZ (aa 123-193, NM_002744.6), the C-terminal C20 domain of human PRKCZ (aa 405-646), or human SET EMD domain (aa 70-226, NM_001122821.2) into pEGFP-N1 or pmRFP-N1 using restriction sites NheI and XhoI, or into pEGFP-C1 using XhoI and KpnI sites to create C- or N-terminally tagged proteins, respectively. The C-KSR-GS construct replacing the vector-based linker (LELKLRILQS) by a glycine-serine rich linker (GGSSGGGGA) was generated by site directed mutagenesis (Q5 Site*-*Directed Mutagenesis Kit, NEB, USA) of the C-KSR-EGFP plasmid. The tandem 2x-C-KSR probe was made by subcloning a second CA3 domain together with a flexible linker sequence (GGSSGGGGA) introduced in the primer in between the two CA3 domains into C-KSR-EGFP using XhoI and KpnI sites. The sequence encoding the nuclear export signal (NES) from mammalian MAPKK kinase (LQKKLEELEL) was inserted by site directed mutagenesis after the start codons of human PRKCZ C20(405-646) or SET EMD(70- 226) to yield plasmids containing C-NES-C20-PRKCZ-EGFP and C-NES-SET-EMD-EGFP, respectively. Two point mutations in the CA3 domain of KSR1 (C359S, C362S) with substitution of the two conserved cysteines with serine residues was introduced using the Q5 Site*-*Directed Mutagenesis Kit. For protein purification from *E.coli,* the KSR1 CA3 domain was cloned into SfiI restriction sites of the pETM11-SUMO3-sfGFP (Saliba et al.) to make pETM11- SUMO3-sfGFP-KSR1. The sequence encoding C-KSR was subcloned into pETM11-SUMO3 via Gibson assembly using Gibson Assembly Cloning Kit (NEB, USA). Sequences encoding N-KSR or C-KSR were subcloned into SfiI sites of pSBbi-pur-H-2Kb (# 111623, Addgene, USA) replacing an existing H-2kb fragment. Primers used are listed in Supplementary Table S3.

### Cell culture, transfection, and stable cell line establishment

Wild-type mouse embryonic fibroblasts (MEF, ATCC CRL-2991), HeLa cells (ECACC, 93021013, a gift from N. Demaurex), and HEK 293T (HEK, ATCC-CRL-11268, a gift from N. Demaurex) were cultured in DMEM (22320, Thermo), MEM (41090, Thermo) and DMEM glutaMAX (31966, Thermo) respectively, containing 10% fetal bovine serum (FBS, Thermo), 1% pen/strep (10000 U/ml penicillin, 10 mg/ml streptomycin, PAN Biotech, Germany). JAWS II (JAWS, ATCC CRL-11904) were grown in alpha minimum essential medium with ribonucleosides, deoxyribonucleosides (Gibco 22571), supplemented with 4 mM L-glutamine (Gibco 25030), 1 mM sodium pyruvate (Gibco 58636), 5 ng/ml GM-CSF (Peprotech, 315-03) 20% FBS, 1% pen/strep. Delipidated FBS was prepared by incubating FBS with 0.5 g/ml lipid removal beads (13358-U, Sigma/Merck) at 4°C overnight with rotation followed by centrifugation (27,000g, 20 min, 4°C) and supernatant filtration using 0.2 μm PES filters (Nalgene). All cell lines were cultured at 37°C, 5% CO2, and passaged 1-2 times per week, and used between passages 10 and 40. All cell lines were tested every 6-12 months either by PCR (LookOut kit, Sigma/Merck) or Mycostrips (Invivogen, ep-mysnc-50) and were negative. Cells were transfected upon reaching 60-70% confluency 2-3 days after seeding using Lipofectamine 2000 (Thermo) in full culture medium without antibiotics for 4-6h. To prepare DNA-lipofectamine complexes, equal volumes of serum- and antibiotic-free media containing DNA or Lipofectamine were mixed and incubated at room temperature for 1h. To obtain stable cell lines via transposition, MEF cells were first co-transfected with pSBbi-pur plasmids encoding the protein of interest and pCMV-(CAT)T7-SB100 plasmid (#34879, Addgene, USA) encoding Sleeping Beauty transposase SB100x. Puromycin (10 μg/mL) selection was performed 2 days after co-transfection. Stable cell lines (N-KSR and C-KSR) were maintained in full culture medium containing puromycin (10 µg/ml). JAWS transfection was achieved using 20 μg DNA and 2 x 10^6^ million cells per 100 μl nucleofection solution from the Amaxa Nucleofector Mouse Dendritic Cell kit (Lonza), and the Y-001 program. Cells were diluted in 400 μl RPMI 1640 Glutamax medium, 25 mM HEPES medium (Thermo, 72400021) immediately after electroporation, incubated for 10 min at RT, then centrifuged at 200 g, 5 min, RT and resuspended in warm full medium and incubated overnight before imaging.

### Immunofluorescence

MEF and HeLa cells seeded on 0.17 mm glass coverslips (Carl Roth) were fixed with 4% paraformaldehyde (PFA, AlfaAesar) in PBS for 30 min at RT followed by 3X washes in PBS. JAWS were fixed in 1%PFA for 10 min. For staining nuclei, fixed cells were incubated with 10 µg/ml Hoechst 33342 (Thermo) for 5 min at followed by three washes with PBS and mounting. For immunostainings, cells were permeabilized with 0.1% Triton X-100 (in PBS) for 5 min at RT, washed 3X by PBS, quenched with 25 mM Glycine/PBS for 10 min at RT, treated with Image-IT-FX (Thermo) for 30 min followed by blocking with 2% bovine serum albumin (BSA)/PBS for 30 min. After that the cells were incubated with primary antibodies at 4°C overnight and with a secondary antibody for 1h in dark humid chambers, antibodies diluted in blocking buffer. 3x washes in PBS were carried out between each step in the procedure. Finally, the coverslips were mounted onto glass slides using 5 µl of ProLong Diamond Mounting medium (Thermo) per coverslip. For labeling recycling endosomes, MEF cells were incubated for 30 min at 37°C with 50 µg/ml Alexa555-conjugated human transferrin (Thermo) in MEF full culture medium containing 0.2% (BSA). After 2x washed with PBS, cells were fixed with 3% PFA for 20 min, quenched with 50 mM ammonium chloride for 15 min (RT), followed by 3 washes in PBS and mounting. Images were acquired either using Zeiss LSM700 or LSM800 confocal systems /Plan apochromat 63x/1.4 objectives and Zeiss Zen 2010b version service pack1, or Nipkow Okagawa Nikon spinning disk confocal imaging system Plan Apo 63x/1.4 DICIII objective and Visiview 4.0 software (Visitron Systems).

### Drug treatments

To induce sphingolipid depletion MEF cells were treated with: 0.5 µM myriocin (or methanol vehicle) from *Mycelia sterilia* in full culture medium, with 2.5 µM fumonisin B1 (methanol vehicle) from *Fusarium moniliforme* in culture medium containing 10% FBS delipidated as above for 3 days, or with 20 µM fumonisin B1 in regular cell culture medium overnight. To induce sphingolipid production MEF cells were treated with 0.5 mM BSA-conjugated palmitic acid complex (Cayman Chemical, USA) for 4h at 37°C in delipidated-FBS medium, and in regular medium with 100 µM Genz-123346 (DMSO vehicle) overnight, or 200 µM C6-ceramide (d18:1/6:0) (Avanti/Merck) for 2h (ethanol vehicle). To induce DAG accumulation cells were treated with 10 µM ionomycin in regular medium for 10 min. To release ceramides locally at the plasma membrane cells were treated with 0.5 U/ml sphingomyelinase from *Staphylococcus aureus* for 30 min in DMEM medium without FBS and antibiotics. After treatment cells were either placed on ice, scraped and used for lipid extraction or fixed with 4% PFA and mounted for microscopy. Alternatively, after SMase treatment, cells were stained with a sphingomyelin probe EGFP-Lysenin (Kiyokawa et al.) (purified from *E.coli,* see below) at 20 µg/ml in serum-free medium for 30 min at 37°C followed by 2 washed in PBS and fixation as described above.

### Image analysis

Image analysis was performed using ImageJ v1.52 (Schneider et al.) NIH. Colocalization analysis was conducted using the JACoP v2.0 plugin (Bolte and Cordelières) that returned Mander’s coefficients for each region of interest (ROI). Mean fluorescence intensities were measured from background-subtracted images. Analyses determining the percentage of cells displaying plasma membrane localization were conducted on images randomized by the Filename_Randomizer.txt macro https://imagej.nih.gov/ij/macros/Filename_Randomizer.txt, by a blinded experimenter that counted the number of plasma membrane-positive cells and total cells in each image using the Cell Counter plugin. For whole-cell fluorescence intensity analyses, confocal z-stacks of 9 frames spaced at 0.5 µm intervals were acquired and sum projections of stacks were created using the Z-Projection tool. Cell and background ROIs were drawn as above on the sum projections images to obtain mean total cellular fluorescence intensities. To eliminate differences between images acquired on LSM700 vs LSM800 microscopes a normalization factor was calculated from images taken on both microscopes from the same coverslip and applied to obtain normalized mean total cellular fluorescence intensities (nMFI).

### Immunoblotting

Cells grown to 80-90% confluency on 6-cm dishes were washed 3x with PBS, scraped off, pelleted, and lysed in RIPA buffer (Thermo 89900) with protease inhibitors (Halt Protease Inhibitor cocktail, 87785) for 30 min at 4°C. Cell debris was pelleted at max speed for 10 min (4°C), and the supernatant was used for loading into the gel. Protein concentration was quantitated using RotiQuant kit (Carl Roth). Lysates were diluted in the SDS Loading Buffer (Roti-Load, Carl Roth) and heated at 95°C for 5 min. 15-30 μg protein were loaded and separated by SDS-PAGE in 4–12% SurePAGE 12-well pre-cast gels (Genscript). Proteins were transferred onto PVDF membranes using iBlot or iBlot2 system (Thermo). After transfer, membranes were blocked in 5%milk/TBS-1%Tween-20 (TBST) for 1h (RT, agitation) followed by incubation with primary antibodies diluted in 3%-milk/TBST overnight at 4°C and 1h incubation at RT with HRP-conjugated secondary antibodies 3%-milk/TBST with multiple washes in PBS between incubations with primary and secondary antibodies. Signals were detected using Immobilon HRP substrate (Merck) on the ImageQuant LAS 4000 mini imaging system (GE, USA).

### Lipid extraction and lipidomics

For lipidomics experiments, lipids were extracted following a modified MTBE protocol (Guri et al.). MEF cells grown to confluency in lipid-free or full culture medium on 6-cm dishes were washed in cold PBS and scraped off in 1 ml PBS. 900 µl and 100 µl cell suspension were transferred to two separate tubes and the cells were pelleted at 400g for 5 min (4°C). The pellets were kept at −80°C or used for protein quantitation (100 µl sample) by resuspension in RIPA buffer, using RotiQuant as above, or for lipid extraction (900 µl sample) as follows. First, pellets were resuspended in 100 µl LC-MS grade water, then 360 µl methanol and the lipid standard mix (see Reagents section) were added, and the tubes vigorously vortexed. Next, 1.2 ml MTBE was added, and the samples were incubated for 1h at RT with shaking. 200 µl of water was added to induce phase separation; samples were incubated at RT for 10 min followed by centrifugation at 1000g, 10 min (RT). The upper organic phase was transferred to 13mm glass tube, 400 µl of the artificial upper phase (MTBE/methanol/water (10:3:1.5, v/v)) was added, the phase separation was repeated, and the collected second upper phase was combined with the first one. The total lipid extract was divided into 2 equal parts: one for sphingolipids (SL) and one for glycerophospholipids (TL). Extracts were dried in a speed vacuum concentrator. After drying, the samples were flushed with nitrogen unless used for further steps. The sphingolipid aliquot was treated with methylamine to remove phospholipids by de-acylation (Clarke method (Clarke and Dawson)). 0.5 mL monomethylamine reagent (MeOH/H2O/n-butanol/Methylamine solution (4:3:1:5 v/v) was added to the dried lipids, tubes sonicated for 6 min, vortexed, followed by the incubation for 1h at 53°C. Finally, the methylamine-treated fraction was dried (as mentioned above). The monomethylamine-treated lipids were subjected to desalting via extraction with *n*-butanol. In short, 300 µl water-saturated *n*-butanol (*n*-butanol/water 2:1 v/v, mixed and prepared at least 1 day before use) was added to the dried lipids. The samples were vortexed and sonicated for 6 min, and 150 µl LC-MS grade water was added. The mixture was vortexed thoroughly and centrifuged at 3200 x g for 10 min. The upper phase was transferred to a 2 mL amber vial. The lower phase extraction was repeated as described above twice more, and the organic phases were combined and dried.

Targeted lipidomic analysis by mass spectrometry was performed according to prior protocols (Jiménez-Rojo et al.) as follows: TL and SL aliquots were resuspended either in 250 µl Chloroform: Methanol (LC-MS grade, 1:1 v/v) or directly in the sample solvent ((chloroform/methanol/water (2:7:1 v/v) + 5mM ammonium acetate). After sonication for 6 min (RT), lipid mixtures were transferred into wells of a 96-well Twin tech PCR plate (Eppendorf) using Hamilton syringes. Sphingolipid aliquots were diluted 1:4, and total lipids were diluted 1:10 in the sample solvent. PCR plate was sealed with EasyPierce foil (Thermo), and the samples were injected into the TSQ Vantage Triple Stage Quadrupole mass spectrometer (Thermo Fisher Scientific) equipped with a robotic sample handler and nanoflow ion source Nanomate HD (Advion Biosciences, USA). The collision energy was optimized for each lipid class. Lipid detection parameters are listed in Supplementary Table S4. Samples were injected at a gas pressure of 30 psi with a spray voltage of 1.2 kV and run on the mass spectrometer operated with a spray voltage of 3.5 kV for positive mode and 3.0 kV for negative mode and a capillary temperature set at 190oC. Sphingolipid and glycerophospholipid species were identified and quantified using multiple reaction monitoring (MRM). Ceramide species were also quantified with a loss of water in the first quadrupole. Each biological replicate was measured twice (2 technical replicates, TR). Each TR consisted of 3 measurements for each transition. Each lipid species was quantified using standard curves created from internal standards. Data analysis was performed using R version 3.6.2 (R Foundation for Statistical Computing). The values where then normalized to the total lipid content of each corresponding lipid extract (mol% of the sum of total PC, PE, PI, PS), except for samples of phagocytosing cells. In this case, because large changes in PC under the phagocytosing condition skewed the normalization, lipid concentrations were normalized first to total protein of each corresponding individual sample, then to the average of the sum of total PC, PE, PI, PS for the entire sample group (control and phagocytosing) processed the same day. Lipid values are thus expressed as mol% of the daily average total lipid mass (mol% of ave tot). Values greater than 100% thus represent values greater than the average total lipid.

### Protein purification from E.coli

Rosetta 2(DE3) *E.coli* cells (Novagen) were transformed with the plasmids expressing His6-SUMO3 tagged KSR probes or GFP alone (control): pETM11-His6-SUMO3- -sfGFP-N- KSR, pETM11-His6-SUMO3-C-KSR-sfGFP, pETM11-His6-SUMO3-sfGFP. Transformed bacteria were grown at 37°C in 1L LB medium containing 100 µg/ml kanamycin, 34 µg/ml chloramphenicol until OD600 0.6-0.8. Protein expression was induced by the overnight incubation with 0.4 mM isopropyl β-D-thiogalactoside (IPTG) at 16°C. The next day bacteria were lysed using the Microfluidizer LM20 high-pressure homogenizer (15,000 psi pressure, 2 cycles) in lysis buffer (50 mM Tris pH 7.5, 500 mM NaCl, 20 mM imidazole, protease inhibitors (1 tablet/50 mL), 0.5 mM DTT) and the soluble fraction was obtained by centrifugation at 16,000g for 1h (4°C). Proteins harboring the His6 tag were captured by incubation with Ni-NTA resin (Qiagen) (2ml resin per bacterial lysate from 1L culture) for 2h, 4°C, 30 rpm rotation. Agarose resin with bound proteins was centrifuged at 1200 rpm, 8 min (4°C), and the slurry was transferred to a column with a cotton filter. The column was washed with 20 ml Lysis buffer, His6-tagged proteins were eluted with 6 ml Elution Buffer (50 mM Tris pH 7.5, 250 mM NaCl, 500 mM imidazole), DTT was added to 5 mM to prevent protein aggregation. Eluted proteins were dialyzed against 50 mM Tris pH 7.5, 200 mM NaCl, 1 mM DTT overnight at 4°C. The His6-SUMO3 tag was cleaved off the proteins by incubation with His-tagged SenP2 protease (EMBL Protein Expression and Purification Core Facility European, Germany) at a molar ratio of SenP2 to protein 40:1. The undigested proteins and the protease were removed using Ni-NTA resin, purified proteins were eluted by applying Elution Buffer 2 (50 mM Tris pH 7.5, 250 mM NaCl, 10 mM imidazole) to Ni-NTA-packed column. The quality of purification was assessed by SDS-PAGE followed by Coomassie staining (RotiBlue kit, CarlRoth). sfGFP-N-KSR was further purified by size exclusion chromatography on the S75 column pre-equilibrated with 50 mM mM Tris, pH 7.4 and 250 mM NaCl.

### Cell lysate preparation from HEK cells

HEK cells grown on 10-cm dishes in HEK full culture medium to 70-80% confluency were transfected as above with plasmids encoding EGFP (control), KSR lipid probes, or PKC- C1(2)-GFP, where 20 µg plasmid DNA/50 µl lipofectamine 2000 was used per dish. After overnight incubation, the cells were trypsinized for 1 min (RT) and resuspended in PBS containing 0.5 mM EDTA. Cells suspension was centrifuged 200g, 5 min (4°C), followed by a wash in PBS. After pelleting, cells were resuspended in 120 µl hypotonic lysis buffer (10 mM Tris-HCl, pH=7.5, Halt protease and phosphatase inhibitors) and incubated on ice for 20 min. Cells were lysed by passing the suspension 15 times through a 30G syringe needle. Protein concentration was measured using a NanoDrop (Thermo). Finally, NaCl and DTT were added to 150 mM and 5 mM, respectively.

### Liposome Microarray-based assay (LiMA)

Liposome-based protein-lipid interaction assay was performed following the protocol described in (Saliba et al.). Lipid mixtures of different compositions were prepared in chloroform: methanol: water (20:9:1 v/v/v). DOPC-based liposomes contained: 0.1 mol % fluorescent lipid PE-Atto647, 0.5% mol stabilizing lipid PE-PEG350, and 2%, 5% or 10% of bovine ceramide mixture (CerMix), C18-ceramide (C18Cer), C18-dihydroceramide (DHCer), sphingomyelin (SM), glucosyl-ceramide (GlcCer), sphingosine (Sph), sphingosine-1- phosphate (S1P), ceramide-1-phosphate (Cer1P), diacylglycerol (DAG), or 10% phosphatidylinositol-4,5-bisphosphate (PI(4,5)P2) (see also Supplementary Table S1). IPM mimicking liposomes were based on a POPC carrier lipid backbone with 0.1 mol % fluorescent lipid PE-Atto647, 0.5% mol stabilizing lipid PE-PEG350, 33% cholesterol, 21.2% DOPS, 15%POPE, and 2%, 5%, or 10% CerMix, C18DHCer, C18Cer, DAG, POPA, SM and Sph. Mixtures containing 5% and 10% GlcCer or 10% DAG did not form liposome, and all liposomes containing DOPA were autofluorescent, thus these spots were excluded from the analysis. Lipids were spotted onto thin agarose layers covered by the protective membrane on top of a glass slide using automated lipid spotting under an inert argon atmosphere (Automatic TLC-spotter4, Camag, Switzerland). A microfluidic chamber with four channels was manufactured from polydimethylsiloxane (PDMS), as described in (Saliba et al.). Then the agarose layers with spotted lipids were glued onto PDMS chambers using a 2:1 mixture of silicone elastomer: curing agent diluted 3× (v/v) in hexane as an adhesive, so the resulting microfluidic chamber contained four channels of 30 spots, 120 lipid spots per chamber. Liposomes were generated by manually injecting the liposome buffer (10 mM HEPES, 150 mM NaCl) into each channel with a Hamilton syringe. Then, 40 µl of the solution containing purified proteins at 5-10 µM or HEK cell lysates in liposome buffer with 5 mM DTT were injected into channels using an automated injection system with a flow rate of 8 µl/min. After incubation of freshly formed liposomes with the GFP-tagged proteins for 1h at RT, the channels were manually washed with 80 µl of liposome buffer, and all spots in the chamber were imaged with a Zeiss Axio Observer Z1 video time-lapse system. All images were taken with the 20x objective; liposomes were imaged with the Cy5 filter set (5 and 10 ms exposure), and GFP-tagged proteins were imaged at different exposure times (1, 5, 10, 25, 50, 75, 100, 150, 200, and 300 ms).

Images were analyzed using CellProfiler version 4.2.5 software (Broad Institute) using a custom pipeline described in detail in (Saliba et al.). A custom-made R script was used to calculate the normalized binding intensity value (NBI) for each protein-lipid interaction. NBI represents a mean of the ratio of GFP to Atto647 fluorescence divided by exposure time at all exposure times for which spots were successfully segmented and is proportional to the amount of GFP-tagged interaction protein recruited to the liposome surface. NBI scores were then expressed as the Log2 ratio of liposome of interest to control liposome containing carrier, fluorescent and stabilizer lipids only.

### Preparation of phagocytic beads and phagocytosis

3 μm carboxyl polystyrene beads (Spherotech, CP30-10) were covalently coupled with purified human IgG (hIgG, Dunn Labortechnik, Germany) by washing 3 times in sterile PBS with centrifugation at maximum speed (18,000 g) at 4°C, activating with 50 mM 1-ethyl-3-(3-dimethylaminopropyl) carbodiimide hydrochloride (EDC-HCl, Carl Roth/2156.1) for 15 min in PBS at room temperature (RT) with rigorous shaking, followed by 3 washes at 4°C in 0.1 M Na2B4O7 buffer (pH 8.0). Finally, 2 µg/µl of hIgG (and 15 ng/µl iFluor647-amine (AAT Bioquest), for fluorescent beads) were added to the beads and incubated overnight at 4°C on a shaker. The next day, the beads were washed 2x with 250 mM glycine/PBS followed by two washes in PBS. To render MEFs phagocytic, cells were transfected with 1 µg/2×10^5^ cells pcDNA3-myc-FCGR2A (Fc receptor) plus KSR1-based probes and where indicated TagRFP-KDEL, mCherry-STIM1 or mCherry-Sec22b at 0.3 µg per 2×10^5^ cells. Cells were washed in full medium 4-6h after transfection. The following day beads were added to the cells at a bead-to-cell ratio 10:1, cells were centrifuged at 200g, 2 min, RT, incubated with the beads for the indicated times at 37°C, washed with PBS, fixed with 4% PFA/PBS for 30 min followed by 3x washes with PBS and mounting as described above.

## Statistics and reproducibility

All statistical analyses were done using Prism (GraphPad Software). P values that are significant are shown above the bars. Two-tailed unpaired t-test were performed when comparing two samples, Welch’s correction was used if F test showed a significant difference in variance. For three or more sample comparisons a one-way ANOVA was employed with in case of negative F test, and Brown-Forsythe and Welch test in case F test showed significant difference in variance, with Dunnett’s post-hoc test. For lipidomic data multiple ratio paired t-test was performed with a two-stage step-up method (Benjamini, Krieger, and Yekutieli), with a false discovery rate set at 5%. Categorical quantification of plasma membrane recruitment was performed by a blinded experimenter on randomized images. Contingency tables were analysed using a Chi-squared test. Cells exhibiting abnormal morphology (multinucleated, high vacuolation) or signs of death (blebbing) were excluded from analysis. A minimum of 5 replicate fields per coverslip are imaged for analysis of fixed cells. These may contain from 1 to tens of cells or 1 to tens of phagosomes which are considered as replicate values. For liposome analysis spots with large aggregrates were excluded from the analysis.

## Data availability

Full datasets, including all microscopy and Western blot source images will be made available on the University of Geneva’s FAIR-compliant data repository Yareta under a CC BY 4.0 licence upon publication. Plasmids generated in this study will be deposited on Addgene upon publication, to be distributed subject to a standard Material’s Transfer Agreement.

## Acknowledgements

We are grateful to the bioimaging and proteins and peptides core facilities at the University of Geneva Medical Centre, Faculty of Medicine as well as the mass spectrometry core facility at the Faculty of Science for their invaluable help. Thanks to Drs. Nicolas Demaurex, Maud Frieden, Brenda Kwak and Amado Carreras-Sureda for their valuable advice. This work was funded by a Professor Dr. Max Cloëtta Foundation Medical Researcher Grant (to PNH), a Swiss National Science Foundation grant (310030_189094, to PNH), a Schmidheiney Foundation subsidy (2022) to PNH. HR acknowledges the Swiss National Science Foundation (310030_184949), the NCCR Chemical Biology (51NF40-185898) and the Leducq Foundation. ACG and LVE acknowledge the financial support of the Louis-Jeantet Foundation, Switzerland.

## Author Contributions

VG designed and performed experiments, analysed data and wrote the manuscript. LVE, IDQ, MA and MCC designed and performed experiments and analysed data. IR, and MM performed experiments and analysed data. HR and ACG designed experiments, provided guidance on analysis and interpretation of data as well as materials. PNH conceived the project, designed and performed experiments and wrote the manuscript.

