## Supplementary_Materials for "Development of genetically-encoded fluorescent KSR1-based probes to track ceramides during phagocytosis"

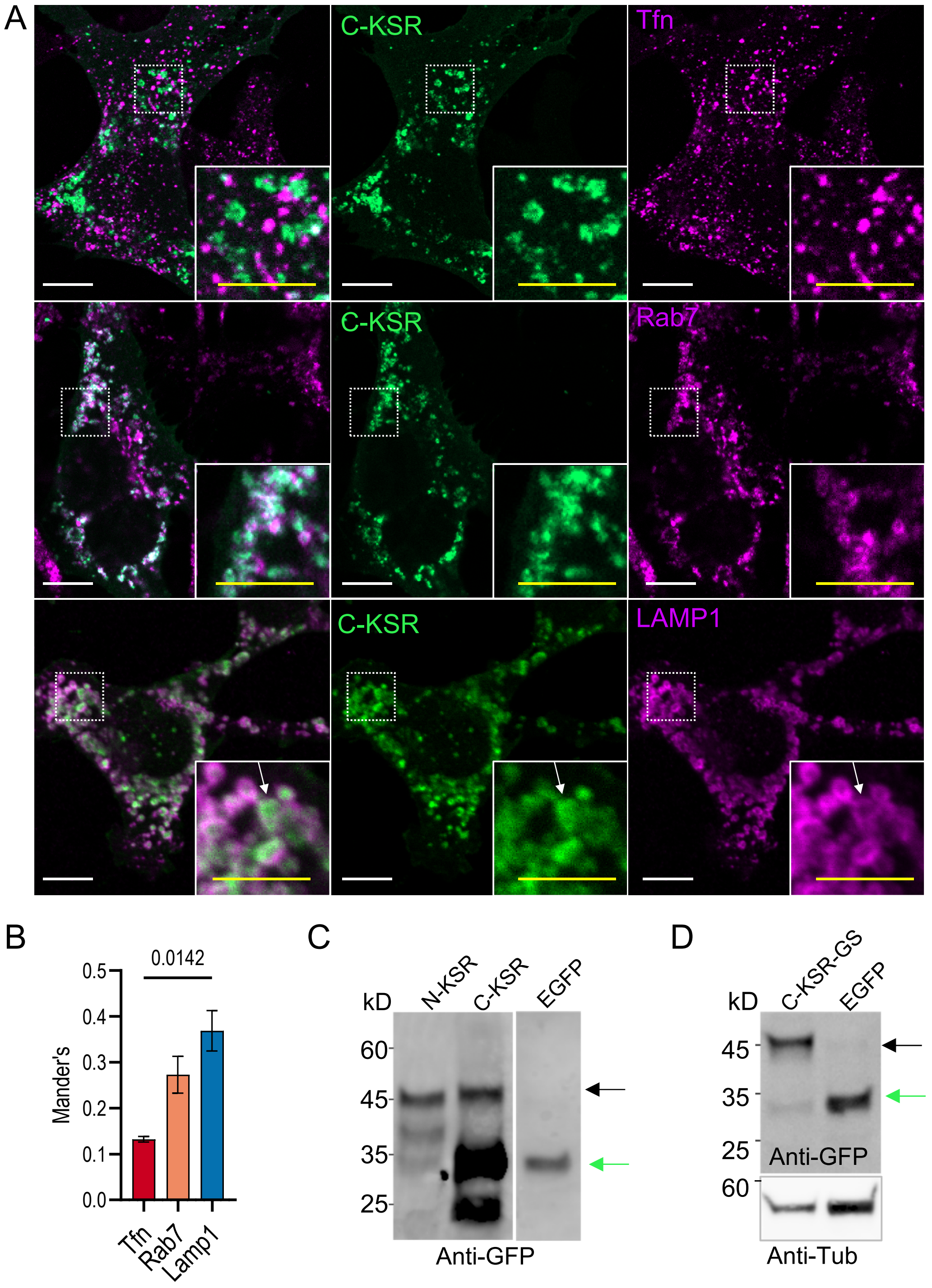


**Figure S1.** *C-KSR localizes to late endosomes and lysosomes.* **A)** Confocal images of MEF cells transfected with C-KSR (green) and loaded for 30 min with recycling endosome marker Alexa555-conjugated transferrin (Tfn, 50 µg/ml, magenta top); or immunostained with late endosome marker anti-Rab7 (magenta middle), or lysosomal marker anti-LAMP1 (magenta bottom). The arrow points to an enlarged vesicular structure showing peripheral C-KSR localization with a hollow lumen, that is surrounded still by LAMP1 staining. **B)** Quantification of co-localization between C-KSR and organelle marker, expressed as Mander's coefficient (n=4/4/14 coverslips for Tfn/Rab7/LAMP1). Bars are means +/- SEM. White bars = 10 µm, yellow bars = 3 µm. **C-D)** Western blot analysis of lysates of MEF cells expressing (EGFP-tagged) N-KSR, C-KSR (C) and C-KSR-GS (D) or soluble cytosolic EGFP control. Immunolabelled with anti-GFP antibodies (and anti-tubulin in D). Black arrow shows the expected size of the intact probe, and the green arrows show the expected size of EGFP alone.

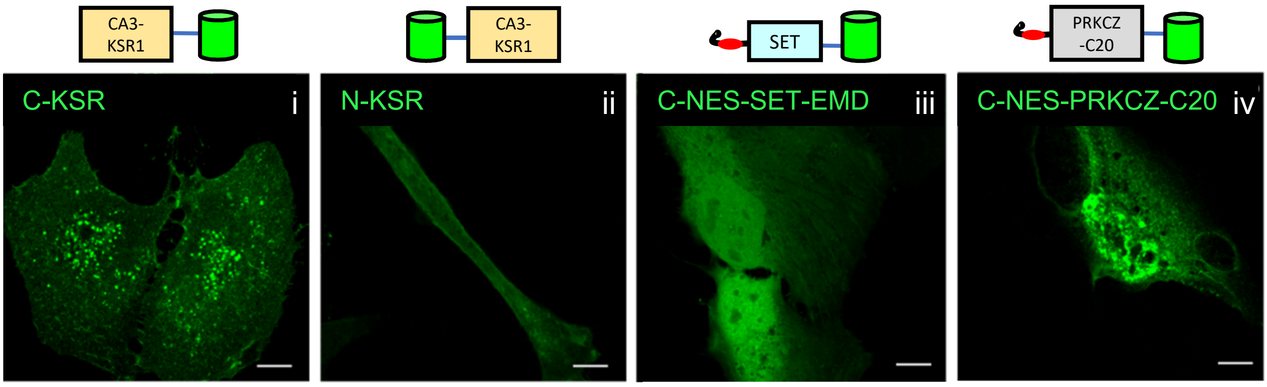


**Figure S2.** *Localization of lipid probe candidates in HeLa cells.* Confocal images of HeLa cells transiently expressing indicated EGFP-tagged proteins (green): C-KSR (i), N-KSR (ii) as well as C-NES-SET-EMD (iii) and C-NES-PRKCZ-C20 (iv) bearing a MAPPKK nuclear export signal (NES). Scale bars, 10 μm.

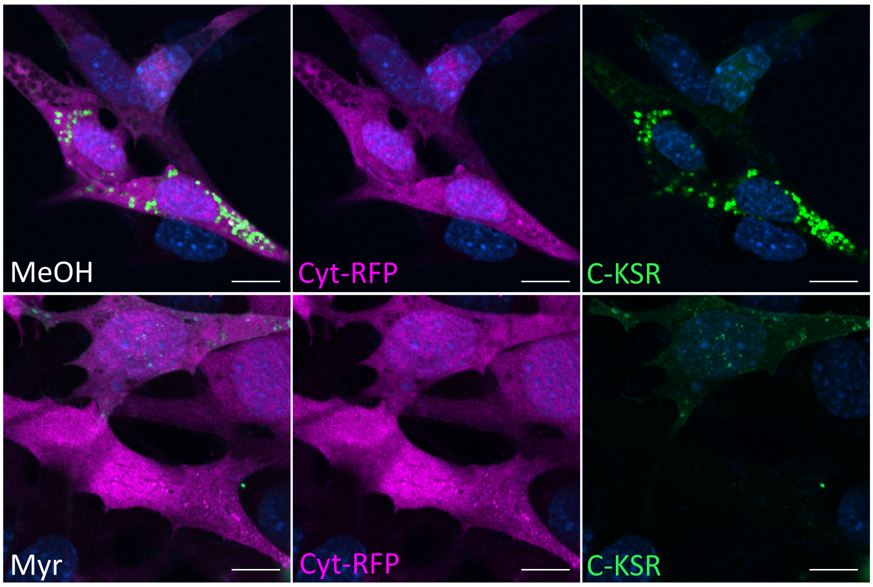


**Figure S3.** *Effect of myriocin on cytosolic RFP*. Confocal images of MEF cells transiently co-transfected with cytosolic TagRFP (Cyt-RFP, magenta) and C-KSR-EGFP (C-KSR, green) and exposed to 0.5 μM myriocin (Myr) for 3d. White bars = 10 μm.

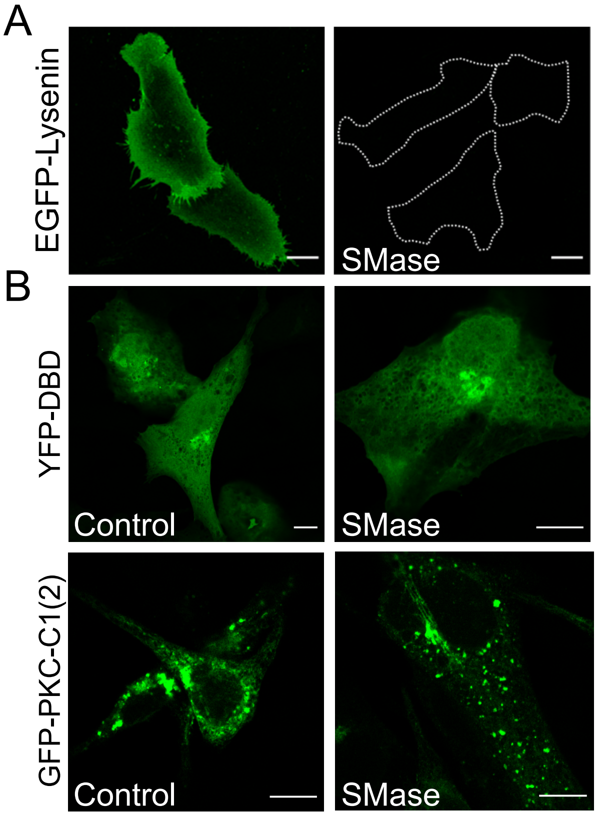


**Figure S4.** *Other potential ceramide probes do not respond to the local ceramide release at the plasma membrane.* **A)** Confocal images of fixed MEF cells pre-treated with or without sphingomyelinase (SMase, 0.5U/ml for 30 min) and stained with a purified sphingomyelin-specific probe EGFP-NT-Lysenin (20 µg/ml). **B)** Confocal images MEF cells transfected with DAG-specific probes YFP-DBD (top) or PKC-C1(2) (bottom), (C), incubated with 0.5 U/ml SMase for 30 min**.** White bars = 10 µm.

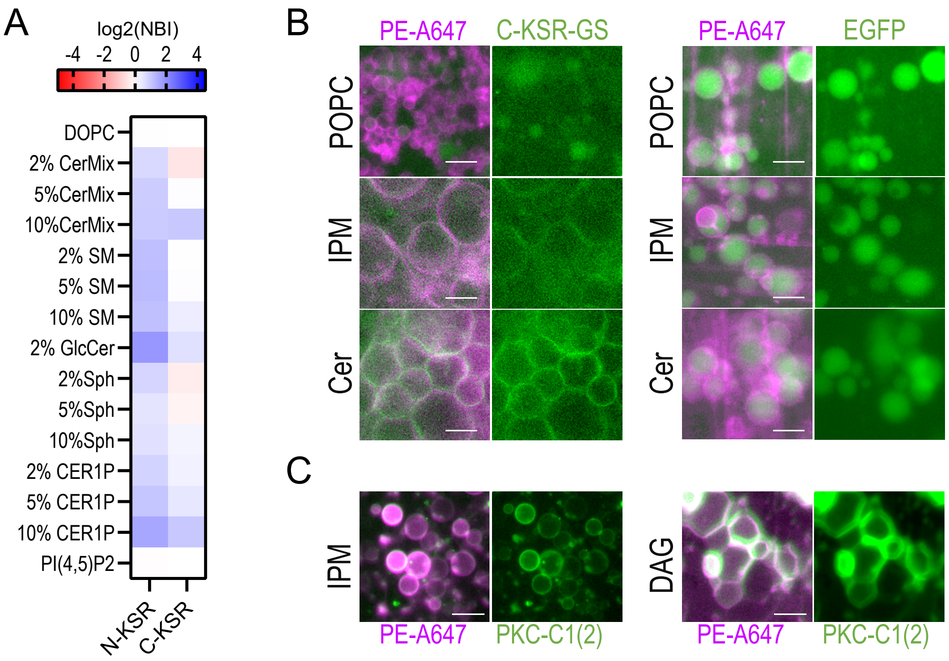


**Figure S5.** *Liposome microarray (LiMA) analysis of KSR1-based probes*. **A)** LiMA heatmap showing average binding of purified N-KSR and C-KSR probes to liposomes based on a DOPC backbone. Binding is expressed as log2 transformed normalized binding intensity (NBI) score ratios, computed to the average of DOPC controls. (See Supplementary Data S2 for full dataset). **B-C)** Representative images of liposomes (containing fluorescent PE-Atto647, magenta) exposed to either C-KSR-GS (left, green), negative control EGFP (right, green) or positive control PKC-C1(2) (C, green). POPC: control liposomes containing only palmitoyl-oleoyl-PC); IPM: inner plasma membrane mimic liposomes (POPC liposomes containing also PE, PS and cholesterol); Cer: IPM liposomes containing also 10%C18-ceramide. DAG: IPM liposomes containing 5% diacylglycerol (DAG). Bar = 10 µm.

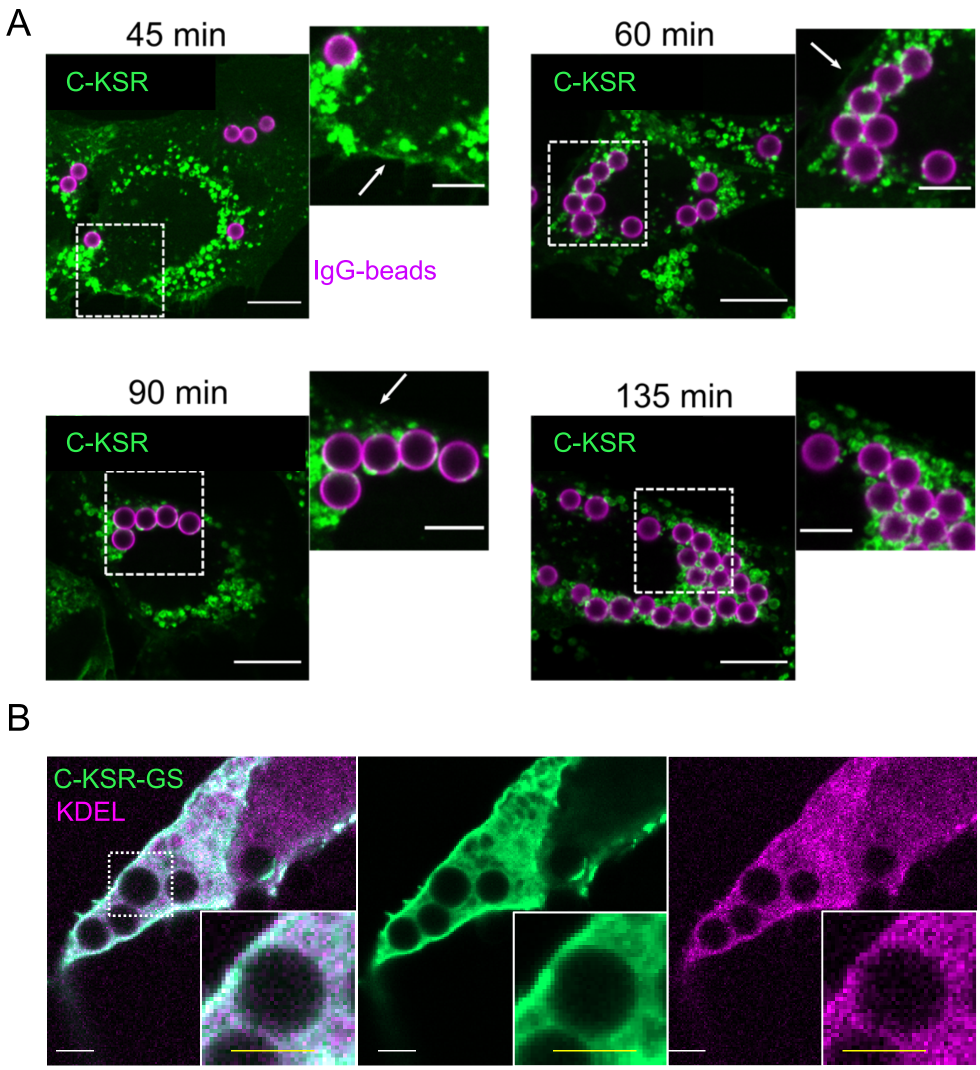


**Figure S6.** *Behavior of KSR probes during phagocytosis*. **A)** Confocal images of phagocytic MEFs expressing C-KSR (green) exposed to IgG-Alexa647-coupled beads at 1:10 cell:bead ratio for 45 (upper left), 60 (upper right), 90 (lower left) and 135 (lower right) min. **C)** Confocal images of phagocytic MEFs co-transfected with C-KSR-GS (green) and ER-targeted soluble TagRFP-KDEL (magenta), and exposed to IgG beads as above for 30 min. White bars = 10 µm, yellow bars = 3 µm.
