## Supplementary_Figures for "Development of genetically-encoded fluorescent KSR1-based probes to track ceramides during phagocytosis"

### Supplementary Materials

**List of Supplementary Materials:**

1. Supplementary Figures:
    - Figures S1 to S6 (file name: Supplementary_Figures.pdf)

2. Supplementary Tables: (below)
    a) **Supplementary Table S1**: List of lipids used in this study.
    b) **Supplementary Table S2**: List of plasmids used and generated in this study.
    c) **Supplementary Table S3**: List of primers used in this study.
    d) **Supplementary Table S4**: Lipid detection parameters for lipidomic analysis

3. Supplementary References: (below)

4. Supplementary Data (separate files):
    a) **Supplementary Data S1**: Lipidomic data of MEF cells treated with myriocin, palmitate, sphingomyelinase, or phagocytosing for 0, 30 and 135 min.
    (file name: Supplementary_Data_S1.xlsx)

    b) **Supplementary Data S2**: Liposome microarray quantification.
    (file name: Supplementary_Data_S2.xlsx)

**Supplementary Table S1.** List of lipids used in liposome microarray analysis.

| Lipid Catalog Name | Lipid Abbreviation | Symbols/Abbreviations/Other names | Source | Catalog# |
|---|---|---|---|---|
| 1,2-dioleoyl-sn-glycero-3-phosphate (sodium salt) | DOPA | DOPA, 18:1 PA, phosphatidic acid | Avanti Polar Lipids | 840875P |
| 1,2-dioleoyl-sn-glycero-3-phospho-(1'-myo-inositol-4',5'-bisphosphate) (ammonium salt) | PI(4,5)P2 | DOPI(4,5)P2, PIP2[4',5'](18:1(9Z)/18:1(9Z)) PI(4,5)P2 | Avanti Polar Lipids | 850155P |
| 1,2-dioleoyl-sn-glycero-3-phosphocholine | DOPC | 18:1 (Δ9-Cis) PC (DOPC) phosphatidylcholine | Avanti Polar Lipids | 850375P |
| 1,2-dioleoyl-sn-glycero-3-phosphoethanolamine | PE-Atto647 | DOPE-Atto647 phosphatidylethanolamine | Atto Tec | AD 647N-161 |
| 1,2-dioleoyl-sn-glycero-3-phosphoethanolamine-N-[methoxy(polyethylene glycol)-350] (ammonium salt) | PE-PEG350 | PE-PEG350 (18:1) phosphatidylethanolamine | Avanti Polar Lipids | 880430O |
| 1,2-dioleoyl-sn-glycero-3-phospho-L-serine (sodium salt) | DOPS | 18:1 PS (DOPS), phosphatidylserine | Avanti Polar Lipids | 840035P |
| 1-2-dioleoyl-sn-glycerol | DAG | 18:1 DG, diacylglycerol | Avanti Polar Lipids | 800811O |
| 1-palmitoyl-2-oleoyl-glycero-3-phosphocholine | POPC | 6:0-18:1 PC (POPC), phosphatidylcholine | Avanti Polar Lipids | 850457P |
| 1-palmitoyl-2-oleoyl-sn-glycero-3-phosphoethanolamine | POPE | 16:0-18:1 PE, POPE, phosphatidylethanolamine | Avanti Polar Lipids | 850757P |
| Ceramide from bovine spinal cord[1] | CER | CerMix, bovine ceramide mixture | Sigma | 22244 |
| cholesterol | Chol | 3β-Hydroxy-5-cholestene, 5-Cholesten-3β-ol, cholesterol | Sigma | C8667 |
| D-erythro-sphingosine | Sph | Sphingosine (d18:1) | Avanti Polar Lipids | 860490P |
| D-erythro-sphingosine-1-phosphate | S1P | Sphingosine-1-Phosphate (d18:1) | Avanti Polar Lipids | 860492P |
| D-glucosyl-ß-1,1'-N-heptadecanoyl-D-erythro-sphingosine | GlcCer | glucosyl-ceramide, C17 Glucosyl(ß) Ceramide (d18:1/17:0) | Avanti Polar Lipids | 860569P |
| N-(hexadecanoyl)-sphing-4-enine-1-phosphocholine | SM | 16:0 SM, C16-sphingomyelin, Egg SM | Avanti Polar Lipids | 860061P |
| N-palmitoyl-ceramide-1-phosphate (ammonium salt) | Cer1P | C1P, C16-ceramide-1-phosphate (d18:1/16:0) | Avanti Polar Lipids | 860533P |
| N-stearoyl-D-erythro-sphinganine | DHCer | C18 Dihydroceramide (d18:0/18:0) | Avanti Polar Lipids | 860627P |
| N-stearoyl-D-erythro-sphingosine | C18Cer | C18 Ceramide (d18:1/18:0) | Avanti Polar Lipids | 860518P |

1. *Made from hydrolysis of total bovine brain sphingomyelins. Major components estimated to be 30% C18-ceramide 35% C24-dihydroceramide in (Vieu et al.)*

**Supplementary Table S1.** List of plasmids used in the study.

| Name (short name) | Vector | Insert | Source |
|---|---|---|---|
| pKSR1-CA3-EGFP (C-KSR) | pEGFP-N1 | Human KSR1 CA3 domain (aa317-400) | This study |
| pKSR1-CA3-mRFP1 (C-KSR) | pmRFP1-N1 | Human KSR1 CA3 domain (aa317-400) | This study |
| pEGFP-KSR1-CA3 (N-KSR) | pEGFP-C1 | Human KSR1 CA3 domain (aa317-400) | This study |
| p2X-KSR1-CA3-EGFP (2x-C-KSR) | pEGFP-N1 | 2x Human KSR1 (aa317-400) fragments in tandem | This study |
| pKSR1-CA3-GS-EGFP (C-KSR-GS) | pKSR1-EGFP | GGSSGGGGA flexible linker between the Human KSR1 CA3 domain (aa317-400) and EGFP | This study |

| | | | |
|---|---|---|---|
| pKSR1-GS-mRFP1 | pKSR1-mRFP1 | GGSSGGGGA flexible linker between the Human KSR1 CA3 domain (aa317-400) and mRFP1 | This study |
| pPRKCZ-C1-EGFP<br>(C-PRKCZ-C1) | pEGFP-N1 | Human PRKCZ C1 domain (aa123-193) | This study |
| pEGFP-PRKCZ-C1<br>(N-PRKCZ-C1) | pEGFP-C1 | Human PRKCZ C1 domain (aa123-193) | This study |
| pPRKCZ-C20-EGFP<br>(C-PRKCZ-C20) | pEGFP-N1 | 20 kD C-terminal domain (C20) of human PRKCZ (aa405-646) | This study |
| pEGFP-PRKCZ-C20<br>(N-PRKCZ-C20) | pEGFP-C1 | 20 kD C-terminal domain (C20) of human PRKCZ (aa405-646) | This study |
| pNES-PRKCZ-C20-EGFP<br>(C-NES-PRKCZ-C20) | pPRKCZ-C20-EGFP | Nuclear export signal from MAPKK preceding the 20 kD C-terminal domain (C20) of human PRKCZ (aa405-646) | This study |
| pSET-EMD-EGFP<br>(C-SET-EMD) | pEGFP-N1 | Human SET earmuff domain (EMD) (aa70-226) | This study |
| pEGFP-SET-EMD<br>(N-SET-EMD) | pEGFP-C1 | Human SET earmuff domain (EMD) (aa70-226) | This study |
| pNES-SET-EMD-EGFP<br>(C-NES-SET-EMD) | pSET-EMD-EGFP | Nuclear export signal from MAPKK preceding the Human SET earmuff domain (EMD) (aa70-226) | This study |
| pETM11-His6-SUMO3-sfGFP | | His6-SUMO3-sfGFP | (Saliba et al.) Gavin laboratory |
| pETM11-SUMO3-EGFP-sfGFP | pETM11-His6-SUMO3-sfGFP | Super-folder-GFP (sfGFP) | (Saliba et al.) Gavin laboratory |
| pETM11-SUMO3-KSR1-CA3-sfGFP<br>(C-KSR) | pETM11-His6-SUMO3-sfGFP | Human KSR1 CA3 domain (aa317-400) | This study |
| pETM11-SUMO3- sfGFP-KSR1-CA3<br>(N-KSR) | pETM11-His6-SUMO3-sfGFP | Human KSR1 CA3 domain (aa317-400) | This study |
| pSBbi-pur H-2Kb | pSBbi-pur | H2Kb (murine MHC-I allele) | Addgene #111623 (Gift from Yewdell laboratory) |
| pCMV(CAT)T7-SB100 | pCMV | SB100X transposase | Addgene #34879 from (Mátés et al.) |
| pSbi-pur-KSR1-CA3-EGFP<br>(C-KSR) | pSBbi-pur | Human KSR1 CA3 domain (aa317-400)-EGFP<br>(C-KSR) | This study |
| pSbi-pur-EGFP-KSR1-CA3<br>(N-KSR) | pSBbi-pur | EGFP-Human KSR1 CA3 domain (aa317-400)<br>(N-KSR) | This study |
| FcgRIIA-cmyc | pcDNA3 | Myc-tagged Fc receptor FcgRIIa (FCGR2A) | (Vieira et al.)<br>Gift from S. Grinstein<br>University of Toronto |
| pGFP-PKC-C1(2)delta | pN2 | GFP-N2-PKCdelta-C1(2) C1(2) domains of PKC delta (rat) | Addgene #21216 (Codazzi et al.) |
| pYFP-DBD | pcDNA3 | C1b diacylglycerol binding domain (DBD) of rat PKC beta II | Addgene #14874 (Gallegos et al.) |
| pCMV6-XL5-mCherry-STIM1 | pCMV6-XL5 | mCherry inserted after the signal sequence of human STIM1 | (Luik et al.)<br>Gift from R. Lewis<br>Stanford University |
| p-mCherry-Sec22b | pCMV-mCherry-C1 | rat Sec22b (ERS24)<br>NM_001025686 | (Petkovic et al.) Gift from T. Galli/ C. Vannier, Institute Jacques Monod Paris |
| pTag-RFP-C | | Cytosolic TagRFP | Evrogen FP141 |

| | | | |
|---|---|---|---|
| pEGFP-N1 | | Vector for C-terminally tagged EGFP constructs | Clontech |
| pEGFP-C1 | | Vector for N-terminally tagged EGFP contructs | Clontech |
| pmRFP-N1 | | Vector for C-terminally tagged mRFP constructs | Addgene plasmid # 54635 Gift from Robert Campbell, Michael Davidson, and Roger Tsien (Campbell et al.) |

**Supplementary Table S3.** List of primers used in the study.

| Primer Name | F/R | Sequence | Purpose |
|---|---|---|---|
| NheI_hKSR1_Ntag_F | For | CTAAGCTAGCGCCACCATGGGGAACCGCATTGATGACG | subcloning hKSR1(aa317-400) into pEGFP-N1 |
| XhoI_hKSR1_Ntag_R | Rev | GATGCTCGAGCCGAGTTAGTGGCAGGAAGG | subcloning hKSR1(aa317-400) into pEGFP-N1 |
| XhoI_hKSR1_Ctag_F | For | TACACTCGAGCTGGCGGTTCCTCTGGTGGTGGTGGTGCGGGGAACCGCATTGATGACG | subcloning hKSR1(aa317-400) into pEGFP-C1 |
| KpnI_hKSR1_Ctag_R | Rev | CACGGGTACCTTACCGAGTTAGTGGCAGGAAGG | subcloning hKSR1(aa317-400) into pEGFP-C1 |
| hKSR_CC359-362SS_F | For | GTGTCCCAGAAGAGCATGATATTTGGAGTGAAG | to introduce mutations (C359S, C362S) into KSR1 |
| hKSR_CC359-362SS_R | Rev | GTGGGAGACCTGCGACAGCCAGGA | to introduce mutations (C359S, C362S) into KSR1 |
| NheI_hPRKCZ_Ntag_F | For | CTAAGCTAGCGCCACCATGAGGAAGCTGTACCGTGCCAAC | subcloning hPRKCZ(aa123-193) into pEGFP-N1 |
| XhoI_hPRKCZ_Ntag_R | Rev | TACACTCGAGAGGCTCTTGGGAAGGCATGAC | subcloning hPRKCZ(aa123-193) into pEGFP-N1 |
| XhoI_hPRKCZ_Ctag_F | For | CTGACTCGAGCTGGCGGTTCCTCTGGTGGTGGTGGTGCGAGGAAGCTGTACCGTGCCAAC | subcloning hPRKCZ(aa123-193) into pEGFP-C1 |
| KpnI_hPRKCZ_Ctag_R | Rev | GCGAGGTACCTTAAGGCTCTTGGGAAGGCATGAC | subcloning hPRKCZ(aa123-193) into pEGFP-C1 |
| NheI_hSET_Ntag_F | For | ATTAGCTAGCGCCACCATGCAGAAGAGGTCAGAATTGATCG | subcloning hSET(aa70-226) into pEGFP-N1 |
| XhoI_hSET_Ntag_R | Rev | GACTCTCGAGATCCATATCGGGAACCAAGTAG | subcloning hSET(aa70-226) into pEGFP-N1 |
| XhoI_hSET_Ctag_F | For | AACGCTCGAGCTGGCGGTTCCTCTGGTGGTGGTGGTGCGCAGAAGAGGTCAGAATTGATCG | subcloning hSET(aa70-226) into pEGFP-C1 |
| KpnI_hSET_Ctag_R | Rev | GACGGGTACCTTAATCCATATCGGGAACCAAGTAG | subcloning hSET(aa70-226) into pEGFP-C1 |
| NheI-hPRKCZ-C20_F | For | TACAGCTAGCGCCACCATGGGTGACACAACGAGCACTTTC | subcloning PRKCZ-C20 (aa 405-646)fragment into pEGFP-N1 |
| XhoI-hPRKCZ-C20_R | Rev | GATGCTCGAGCGACTCCTCGGTGGACAGC | subcloning PRKCZ-C20 (aa 405-646)fragment into pEGFP-N1 |
| XhoI-hPRKCZ-C20_F | For | TACACTCGAGCTGGCGGTTCCTCTGGTGGTGGTGGTGCGGGTGACACAACGAGCACTTTC | subcloning PRKCZ-C20 (aa 405-646)fragment into pEGFP-C1 |
| KpnI-hPRKCZ-C20_R | Rev | CAATGGTACCCGACTCCTCGGTGGACAGC | subcloning PRKCZ-C20 (aa 405-646)fragment into pEGFP-C1 |
| XhoI-hKSR1-2X-F | For | GAACCTCGAGGGGAACCGCATTGATGACG | subcloning hKSR1(aa317-400) into pKSR1-EGFP |
| KpnI-hKSR1-2X-R | Rev | TACAGGTACCGACCGAGTTAGTGGCAGGAAGG | subcloning hKSR1(aa317-400) into pKSR1-EGFP |
| PRKCZ_C-C20-NES-F | For | GAGCTGGATGAGGCACCGGTCGCCACCGTGAGCAAGGGCGAGGAG | addition of MAPKK nuclear export signal to PRKCZ-C20-EGFP |
| PRKCZ_C-C20-NES-R | Rev | AAGCTCTTCCAACTTTTTCTGCAGAGCCATGGTGGCGACCGGTAG | addition of MAPKK nuclear export signal to PRKCZ-C20-EGFP |
| SET_N-C20-NES-F | For | GAGCTGGATGAGGCACCGGTCGCCACCCAGAAGAGGTCAGAATTGATCGCC | addition of MAPKK nuclear export signal to SET-EGFP |
| SET_N-C20-NES-R | Rev | AAGCTCTTCCAACTTTTTCTGCAGAGCCATGGTGGCGCTAGCGGA | addition of MAPKK nuclear export signal to SET-EGFP |

| | | | |
|---|---|---|---|
| pETM11-lin-F | For | ACTGAGATCCGGCTGCTAAC | linearization of vector pETM11-SUMO3-EGFP-sfGFP (Gibson assembly) |
| pETM11-lin-R | Rev | GGATCCACCGGTCTGTTGC | linearization of vector pETM11-SUMO3-EGFP-sfGFP (Gibson assembly) |
| KSR-EGFP-F | For | CAACAGACCGGTGGATCCGGGAACCGCATTGATGACG | KSR1 insert amplification to subclone into linearized pETM11-SUMO3-EGFP-sfGFP via Gibson assembly |
| EGFP-R | Rev | TAGCAGCCGGATCTCAGTTTACTTGTACAGCTCGTCCATG | insert amplification to subclone EGFP-tagged probes into a linearized vector pETM11-SUMO3-EGFP-sfGFP via Gibson assembly |
| SfiI-KSR1-F | For | CATGGCCCACAGGGCCTGGGGAACCGCATTGATGACG | for subcloning KSR1 fragment onto SfiI sites of vector pETM11-SUMO3-SfiI-sfGFP |
| Sfi-KSR1-R | Rev | TACGGCCGATATGGCCTTACCGAGTTAGTGGCAGGAAGG | for subcloning KSR1 fragment onto SfiI sites of vector pETM11-SUMO3-SfiI-sfGFP |
| SfiI-probe-EGFP-F | For | CGTGGCCTCTGAGGCCTGAACCGTCAGATCCGCTAG | subloning C-terminally tagged KSR1 into pSBbi-pur |
| SfiI-probe-EGFP-R | Rev | TACGGCCTGACAGGCCTTACTTGTACAGCTCGTCCATG | subloning C-terminally tagged KSR1 into pSBbi-pur |
| C-KSR-GSplus-mut-F | For | ggtggtggtgcgTCGACGGTACCGCGGGCC | deletes LELKLRILQS linker connecting KSR1 CA3 domain to EGFP (in C-KSR constructs) and adds GGSSGGGGA linker. |
| C-KSR-GSplus-mut-R | Rev | accagaggaaccgccCCGAGTTAGTGGCAGGAAGGATATTCTACAG | deletes LELKLRILQS linker connecting KSR1 CA3 domain to EGFP (in C-KSR constructs) and adds GGSSGGGGA linker. |

**For = forward, Rev = reverse, F/R = forward or reverse*

**Supplementary Table S4.** Lipid detection parameters for lipidomic analysis

| **Lipid class** | **Standard** | **Polarity** | **Mode** | **m/z ion** | **Collision energy eV** |
|---|---|---|---|---|---|
| Phosphatidylcholine $[M+H]^+$ | DLPC | + | Product ion | 184.07 | 30 |
| Phosphatidylethanolamine [M+H]+ | PE31:1 | + | Neutral ion loss | 141.02 | 20 |
| Phosphatidylinositol [M-H]- | PI31:1 | - | Product ion | 241.01 | 44 |
| Phosphatidyserine [M-H]- | PS31:1 | - | Neutral ion loss | 87.03 | 23 |
| Cardiolipin [M-2H]2- | CL56:0 | - | Product ion | Acyl chain | 32 |
| Ceramide | C17Cer | + | Product ion | 264.34 | 25 |
| Hexosylceramide | C8GC | + | Product ion | 264.34 | 30 |
| Sphingomyelin | C12SM | + | Product ion | 184.07 | 26 |
